# Identification and characterization of a human MORC2 DNA binding region that is required for gene silencing

**DOI:** 10.1101/2024.06.05.597643

**Authors:** Nikole L. Fendler, Jimmy Ly, Luisa Welp, Henning Urlaub, Seychelle M. Vos

## Abstract

The eukaryotic microrchidia (MORC) protein family are DNA <u>g</u>yrase, <u>H</u>sp90, histidine <u>k</u>inase, Mut<u>L</u> (GHKL)-type ATPases involved in gene expression regulation and chromatin compaction. The molecular mechanisms underlying these activities are incompletely understood. Here we studied the full-length human MORC2 protein biochemically. We identified a DNA binding site in the C-terminus of the protein, and we observe that this region is heavily phosphorylated in cells. Phosphorylation of MORC2 reduces its affinity for DNA and appears to exclude the protein from the nucleus. We observe that DNA binding by MORC2 reduces its ATPase activity and that MORC2 can topologically entrap multiple DNA substrates between its N-terminal GHKL and C-terminal coiled coil 3 dimerization domains. Finally, we observe that the MORC2 C-terminal DNA binding region is required for gene silencing in cells. Together, our data provide a model to understand how MORC2 engages with DNA substrates to mediate gene silencing.

## Introduction

The microrchidia (MORC) protein family are DNA <u>g</u>yrase, <u>H</u>sp90, histidine <u>k</u>inase, and Mut<u>L</u> (GHKL)-type ATPases involved in gene expression regulation (1). MORC proteins share a common architecture consisting of an N-terminal GHKL domain and a predicted C-terminal dimerization domain (2). Eukaryotic MORC proteins likely contribute to gene silencing by compacting chromatin. They are localized to specific genomic regions by post-translational modifications on themselves or histone proteins, or by association with protein complexes that associate with MORC proteins (3–8).

The exact molecular mechanisms underlying DNA binding and chromatin compaction by MORC family proteins are unknown. Members of the GHL subclass of the GHKL-type ATPase family, including type II topoisomerases, MutL, and Hsp90, possess an N-terminal GHKL domain that dimerizes in the presence of ATP and a second C-terminal dimerization interface. GHL ATPases trap substrates in the lumen formed between the two dimerization interfaces (9–13). MORC proteins could similarly engage DNA between their two dimerization interfaces, which could lead to chromatin compaction if multiple DNA strands are engaged simultaneously (1, 3, 14–16). Indeed, *C. elegans* MORC-1 can topologically entrap DNA substrates, supporting this model (15). The regions of MORC-1 that engage DNA, however, have not been identified.

Humans encode five MORC proteins (MORC1-4, and SMCHD1). MORC2, MORC3, and SMCHD1 are constitutively expressed in all cell types whereas MORC1 and MORC4 expression is limited to specific tissues (17). At a structural level, MORC1/MORC2 and MORC3/MORC4 share a similar domain architecture while SMCHD1 is structurally distinct from the other MORC family members (16, 18–21). MORC proteins appear to regulate specific gene subsets. For example, MORC1 is responsible for silencing transposable elements in the male germline, MORC2 silences retrotransposons and genes with long exons, MORC3 silences virally derived DNAs, and SMCHD1 silences expression of the *Dux4* transcription factor (5, 8, 22–26). To date, the only MORC family protein that has been studied biochemically in its full-length form is *C. elegans* MORC-1 (15). All other biochemical and structural studies of MORC family proteins have been conducted with N-terminal truncations that lack the predicted C-terminal dimerization interface, and thus significant questions regarding MORC association with DNA remain unanswered (16, 18–20).

To study how MORC family proteins could associate with DNA in their full-length form, we have biochemically investigated full-length human MORC2. Human MORC2 is a 1032 amino acid protein containing an N-terminal GHKL ATPase domain bifurcated by a coiled coil, a CW-type zinc finger domain (CW), a predicted chromo-like domain (CD), and two predicted C-terminal coiled coil domains (**Figure 1A**). Human MORC2 represses actively transcribed double strand DNA viruses (27), retroelements, particularly evolutionarily young, long interspersed nuclear element 1 (LINE1) (24, 28–32), and intronless and long-exon containing genes (5, 24). Gene silencing by MORC2 requires its ATPase activity and its three coiled coil domains (5). Although MORC2 contains two possible chromatin binding domains (CW and CD), MORC2’s CW domain does not bind histone tails (33) and the CD is dispensable for MORC2 silencing in cells (5). The MORC2 coiled coil 1 domain is the only region of MORC2 that has been shown to bind DNA (18). This interface lies on the periphery of the protein, and it is unclear how it would promote topological entrapment of DNA substrates.

**Figure 1.**
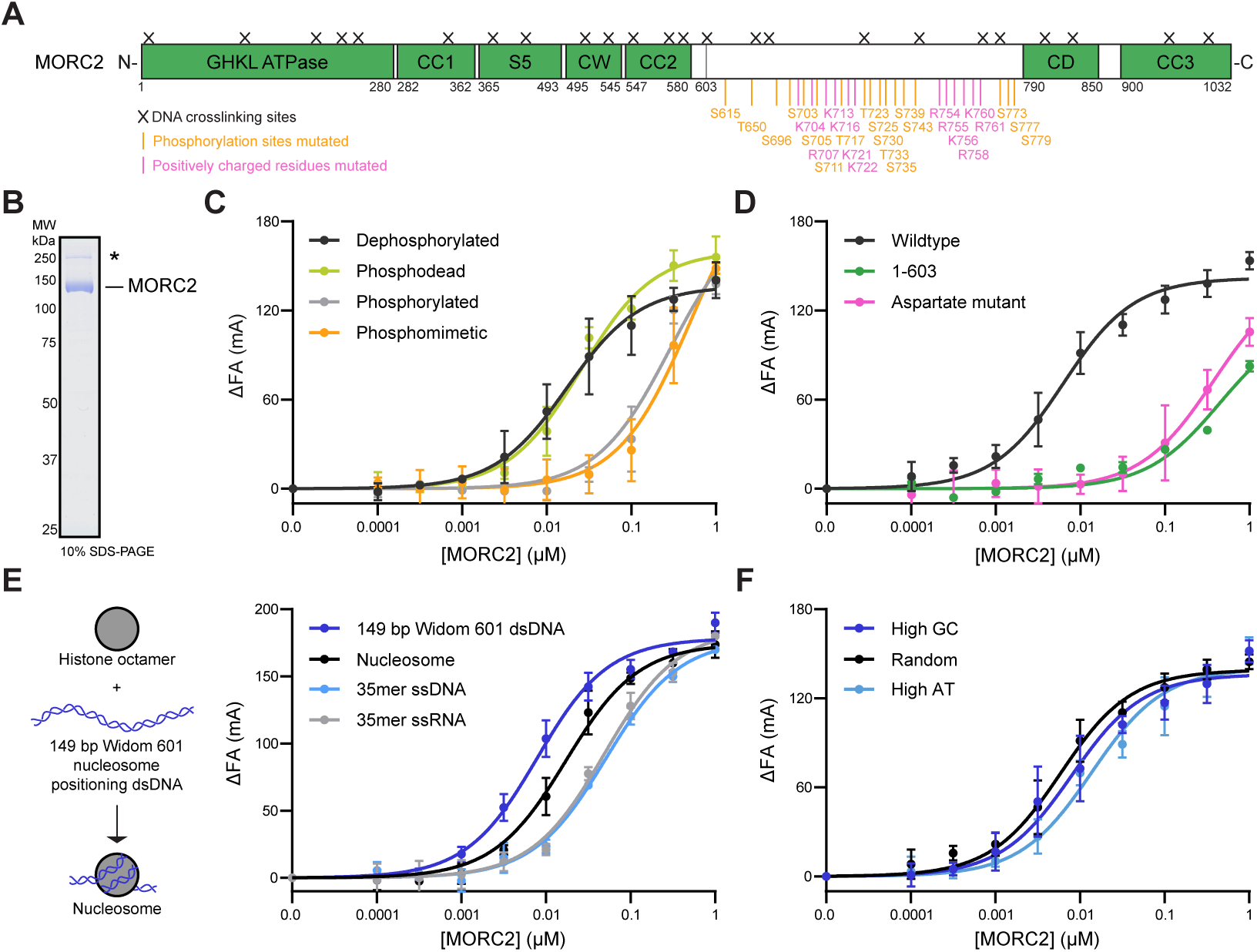
The MORC2 C-terminus contains a DNA binding region that is regulated by phosphorylation. **A.** Domain architecture of human MORC2. Coiled coil domains (CC), S5 transducer domain of the GHKL ATPase module (S5), a CW-type zinc finger domain (CW), and a predicted chromodomain (CD) are indicated. Purified MORC2 was cross-linked to DNA with 1 mM mechlorethamine for 30 minutes at 37°C or cross-linked to DNA by UV light at 254 nm for 10 minutes on ice, and samples were digested by trypsin. Resulting fragments were enriched by TiO_2_ and analyzed by mass spectrometry (**Methods**). Identified amino acid positions cross-linked to DNA are denoted by a black cross. Positively charged residues and phosphorylation sites mutated in this study are noted in pink and orange, respectively, beneath the primary structure. **B.** SDS-PAGE gel of purified human MORC2 protein (3 µg) stained with Coomassie blue stain. Contaminant band marked with an asterisk (**SI Table 1**). **C.** Phosphorylation reduces MORC2 affinity for DNA. Dephosphorylated, phosphorylated, phosphodead, and phosphomimetic MORC2 were assessed for DNA binding using fluorescence anisotropy. MORC2 was titrated with 1 nM FAM-labelled 35 base pair duplex DNA (**Methods**). Binding curves were fit using a quadratic binding equation. Error bars correspond to the standard deviation between three replicate experiments. **D.** Assessment of DNA binding by wildtype, aspartate mutant, and 1-603 MORC2. MORC2 constructs were titrated and incubated with 1 nM FAM-labelled 35 base pair duplex DNA (**Methods**). Binding curves were fit as in 1C. Error bars correspond to the standard deviation between three replicate experiments. **E.** MORC2 nucleic acid binding as measured by fluorescence anisotropy. MORC2 was titrated and incubated with 1 nM 5’ FAM-labelled 149 bp Widom 601 double stranded DNA, nucleosome, 35 nucleotide single stranded RNA, and 35 base pair single stranded DNA (**Methods**). Binding curves were fit as in 1C. Error bars correspond to the standard deviation between three replicate experiments. **F.** MORC2 DNA binding to DNA sequences of different AT/GC content as assessed by fluorescence anisotropy. MORC2 was titrated and incubated with 1 nM 5’ FAM-labelled 35 base pair duplex DNA with a high GC (71% GC content), high AT (31% GC content), or a random sequence with 49% GC content (**Methods**). Binding curves were fit as in 1C. Error bars correspond to the standard deviation between three replicate experiments.

Here we have identified a C-terminal region of human MORC2 that interacts robustly with DNA. The C-terminal DNA binding region is located within the predicted substrate binding lumen of the MORC2 dimer. Phosphorylation within the C-terminal DNA binding region reduces MORC2 association with DNA and results in MORC2 exclusion from the nucleus. Further, DNA binding diminishes MORC2 ATP hydrolysis activity. Finally, we show that MORC2 dimerization at its N- and C-termini results in the selective capture of circular DNA substrates, suggesting that it can encircle DNA and capture multiple DNA fragments simultaneously. Mutations to the MORC2 DNA binding region prevent capture of circular DNA substrates and ablate its gene silencing activity in cells. Together, our work identifies a conserved region of MORC2 that engages DNA substrates and provides a model for understanding how MORC2 and other MORC proteins contribute to gene silencing.

## Material and Methods

### Molecular cloning and protein expression

MORC2 (Uniprot ID: Q9Y6X9, isoform 1) was amplified from K562 cDNA and inserted into insect cell expression vector 438-B (Addgene plasmid: 55219) (N-terminal 6x His (His_6_) tag followed by Tobacco etch virus (TEV) protease cleavage site) and 438-C (Addgene plasmid: 55220) (N-terminal His_6_-Maltose binding protein (MBP) followed by TEV protease cleavage site) by ligation independent cloning (LIC) (34). 438-C MORC2 was used as a template for PCR amplification to introduce mutations and generate MORC2 mutants N39A, E35A, aspartate DNA binding mutant (K704D, R707D, K713D, K716D, K721D, K722D, R754D, R755D, K756D, R758D, K760D, R761D), Subset N (K704D, R707D, K713D, K716D, K721D, K722D), Subset C (R754D, R755D, K756D, R758D, K760D, R761D), alanine DNA binding mutant (K704A, R707A, K713A, K716A, K721A, K722A, R754A, R755A, K756A, R758A, K760A, R761A), 1-603, and phosphodead (S615A, T650A, S696A, S703A, S705A, S711A, T717A, T723A, S725A, S730A, T733A, S735A, S739A, S743A, S773A, S777A, S779A) by sequence-dependent ligation independent cloning (SLIC). For phosphomimetic MORC2 (S615D, T650E, S696D, S703D, S705D, S711D, T717E, T723E, S725D, S730D, T733E, S735D, S739D, S743D, S773D, S777D, S779D), the region between amino acid 696 and 779 with phosphomimetic mutations was generated as a synthetic gene block (Integrated DNA Technologies) and cloned into MORC2 in 438-C by Sequence and Ligation Independent Cloning (SLIC). For coiled coil 3, the region between amino acids 900 and 1032 was amplified by PCR and inserted into *E. coli* expression vector 1B (N-terminal 6x His (His_6_) tag followed by Tobacco etch virus (TEV) protease cleavage site) (Addgene plasmid: 29653).

For PiggyBac EGFP-tagged proteins, we first generated a tetracycline inducible EGFP plasmid containing the puromycin resistance marker (puromycin N-acetyltransferase). EGFP-TEV-S with multiple cloning sites followed by the bGH poly(A) signal amplified by PCR and inserted downstream of the tetracycline-responsive promoter between cut sites NheI and SgfI in the HP138 puro plasmid (Addgene plasmid: 134246). MORC2 constructs were amplified by PCR and inserted downstream of TetO: EGFP-TEV-S between the KpnI and NotI restriction sites to generate tetracycline inducible EGFP-TEV-S-MORC2. These donor plasmids contained puromycin N-acetyltransferase (puromycin resistance), reverse tetracycline-controlled transactivator, and the tetracycline inducible EGFP-tagged transgene flanked by piggyBac inverted terminal repeats. To add the SV40 NLS (PKKKRKV) (35), an oligo duplex containing the SV40 NLS along with 20 bp homology arms were cloned upstream of EGFP-tagged MORC2 constructs cut using the NEB Gibson Assembly Master Mix (E2611L). For spike in mRNA plasmids, Nano and Firefly luciferase flanked with the *Xenopus Laevis* globin 5’ and 3’ UTR was cloned downstream of the T7 promoter using Gibson assembly. All clones were verified by Sanger sequencing and full plasmid sequencing (Primordium or Plasmidsaurus).

Purified plasmid DNA (0.3-0.8 µg) was electroporated into DH10αEMBacY cells to generate bacmids (36). Bacmids were prepared from positive clones by isopropanol precipitation and transfected into Sf9 cells grown in ESF921 (Expression Systems) with X-tremeGENE9 transfection reagent (Sigma) to generate V0 virus. V0 virus was harvested 48– 72 hr after transfection. V1 virus was produced by infecting 25 mL of Sf9 or Sf21 cells grown at 27°C, 300 rpm with 25-150 µL V0 virus. Cells were maintained at 1E6 cells/mL during virus production. V1 viruses were harvested 48-72 hours after proliferation arrest and stored at 4°C. For protein expression, 600 mL of Hi5 cells grown in ESF921 medium were infected with 100-300 µL of V1 virus and grown for 60 hours at 27°C. Cells were maintained at 1E6 cells/mL during expression. Cells were harvested by centrifugation (238xg, 4°C for 30 min), resuspended in lysis buffer at 4°C (500 mM NaCl, 20 mM Na•HEPES pH 7.4, 10% glycerol (v/v), 5 mM beta mercaptoethanol (BME), 30 mM imidazole pH 8.0, 2 μM pepstatin A, 0.7 μM leupeptin, 1 mM phenylmethylsulfonyl fluoride, 2.8 mM benzamidine), snap frozen, and stored at −80°C.

Coiled coil 3 was expressed in Bl21 (DE3) RIL cells. 2xYT media supplemented with 100 µg/mL kanamycin and 34 µg/mL chloramphenicol was inoculated an overnight culture. Cells were grown at 37°C, 160 rpm until reaching an optical density 600 nm 0.4-0.5. Protein expression was induced by adding 0.5 mM of isopropyl β-D-1-thiogalactopyranoside (IPTG) and cells were grown for an additional 4-5 hours at 37°C. Cells were harvested by centrifugation at 4000 rpm, 4°C for 30 min, resuspended in lysis buffer at 4°C, snap frozen in liquid nitrogen, and stored at -80°C.

### Protein purification

All steps were performed at 4°C unless otherwise specified. Frozen cell pellets were thawed in a room temperature water bath, and cells were lysed by sonication. Lysates were clarified by centrifugation. Clarified lysates were filtered through a 5 µm syringe filter, followed by additional filtration through 0.45 µm syringe filters. The filtered lysate was applied to a 5 mL HisTrap (Cytvia) column equilibrated in lysis buffer. The HisTrap column was washed with 10 CV of lysis buffer followed by 4 CV of High Salt buffer (1M NaCl, 20 mM Na•HEPES pH 7.4, 10% glycerol (v/v), 5 mM BME, 30 mM imidazole pH 8.0) followed by 4 CV of lysis buffer, and 4 CV of Low Salt buffer (150 mM NaCl, 20 mM Na•HEPES pH 7.4, 10% glycerol (v/v), 5 mM BME, 30 mM imidazole pH 8.0). Tandem 5 mL HiTrap SP (Cytiva) and 5 mL HiTrap Q HP (Cytiva) columns equilibrated in Low Salt buffer were attached directly to the bottom of the HisTrap column. Protein was eluted from the HisTrap column with 9 CV of Nickel Elution buffer (150 mM NaCl, 20 mM Na•HEPES pH 7.4, 10% glycerol (v/v), 5 mM BME, 500 mM imidazole pH 8.0), after which the HiTrap S and HiTrap Q columns were detached from the HisTrap column. The HiTrap S and HiTrap Q columns were washed with 4 CV of Low Salt buffer and protein was eluted from the HiTrap S and HiTrap Q columns separately by a gradient of 0-100% Low Salt buffer to High Salt buffer (flow rate of 1.5 mL/min for 30 minutes). Peak fractions were analyzed by 10% Tris-glycine SDS-PAGE followed by Coomassie staining. Fractions containing full-length protein were combined with 1.5 mg of His_6_-TEV protease (37) and dialyzed overnight in 7K MWCO SnakeSkin tubing (Fisher) against 1L of Dialysis buffer (500 mM NaCl, 20 mM Na•HEPES pH 7.4, 10% glycerol (v/v), 5 mM BME, 30 mM imidazole pH 8.0, 1 mM MnCl_2_). Dephosphorylated protein was generated by the addition of 0.4 mg of lambda protein phosphatase (prepared in house) at this step.

Protein was removed from the SnakeSkin tubing and applied to a 5 mL HisTrap column equilibrated in Lysis Buffer to remove uncleaved protein, TEV protease, and the tag. Protein was concentrated in an Amicon (Millipore) 15 mL centrifugal concentrator 30K MWCO to 4-5 mL. The protein was applied to a S200 16/600 pg column (Cytvia) equilibrated in SEC buffer (500 mM NaCl, 20 mM Na•HEPES pH 7.4, 10% (v/v) glycerol, and 1 mM tris(2-carboxyethyl)phosphine (TCEP)). Peak fractions were analyzed by 10% Tris-glycine SDS-PAGE followed by Coomassie staining. Peak fractions which contained full-length protein were pooled and concentrated in an Amicon (Millipore) 4 mL centrifugal concentrator 30K MWCO to 50-100 µM, aliquoted, flash frozen, and stored at −80°C. Typical protein preparations yield 1-2 mg of wildtype MORC2 from 1.2 L of insect cell culture. MBP-tagged MORC2 was prepared as described above without the addition of TEV protease in the overnight dialysis and with a Superose 6 10/300 (Cytvia) column for the size exclusion.

### Coiled coil 3 purification

All steps were performed at 4°C unless otherwise specified. Frozen cell pellets were thawed in a room temperature water bath, and cells were lysed by sonication. Lysates were clarified by centrifugation (20,000 rpm, 30 minutes, A27 rotor). The lysate was applied to 2 x 5 mL HisTrap (Cytvia) columns equilibrated in lysis buffer. The HisTrap column was washed with 10 CV of lysis buffer followed by 4 CV of High Salt buffer (1M NaCl, 20 mM Na•HEPES pH 7.4, 10% glycerol (v/v), 5 mM BME, 30 mM imidazole pH 8.0) followed by 4 CV of lysis buffer. Protein was eluted from the HisTrap columns with a gradient from 0-100% Nickel Elution buffer (500 mM NaCl, 20 mM Na•HEPES pH 7.4, 10% glycerol (v/v), 5 mM BME, 500 mM imidazole pH 8.0) with a flow rate of 1.5 mL/min for 30 minutes. Peak fractions were analyzed by 15% Tris-glycine SDS-PAGE stained with Coomassie. Fractions containing full-length protein were combined with 1.5 mg of His_6_-TEV protease (37) and dialyzed overnight in 7K MWCO SnakeSkin tubing (Fisher) against 1L of Dialysis buffer (500 mM NaCl, 20 mM Na•HEPES pH 7.4, 10% glycerol (v/v), 5 mM BME, 30 mM imidazole pH 8.0).

Protein was removed from the SnakeSkin tubing and applied to a 5 mL HisTrap column equilibrated in Lysis Buffer to remove uncleaved protein, TEV protease, and the tag. Protein was concentrated in an Amicon (Millipore) 15 mL centrifugal concentrator 3K MWCO to 4-5 mL. The protein was applied to a S75 16/600 pg column (Cytvia) equilibrated in SEC buffer (500 mM NaCl, 20 mM Na•HEPES pH 7.4, 10% (v/v) glycerol, and 1 mM TCEP). Peak fractions were analyzed by 15% Tris-glycine SDS-PAGE followed by Coomassie staining. Peak fractions which contained full-length protein were pooled and concentrated in an Amicon (Millipore) 4 mL centrifugal concentrator 3K MWCO to 100-200 µM, aliquoted, flash frozen, and stored at −80°C.

### Phospho-mass spectrometry analysis

Protein samples were excised from Tris-glycine SDS-PAGE gels stained with Coomassie blue, processed, and subjected to trypsin digest. Digested samples were separated by liquid chromatography and analyzed by tandem mass spectrometry. MS/MS data were searched against the sequence for human MORC2 and the *Trichoplusia ni* genome using SEQUEST. Phosphorylation modifications were identified computationally from peptide spectra without additional enrichment.

### DNA-protein cross-linking and mass spectrometry

#### MORC2-65mer DNA complex reconstitution, crosslinking, and sample processing

3.3 µL MORC2 protein (132.3 µM, corresponding to 436.6 pmol, 51.6 µg protein) was incubated with 5.7µL 65mer DNA (100 µM, corresponding to 570 pmol) for 10 min, on ice. The sequence of the 65 base pair DNA is as follows: 5’-/6-FAM-ATT CTC CAG GGC GGC CGC GTA TAG GGT CCA TCA GAA TTC GGA TGA ACT CGG TGT GAA GAA AGA TC-3’

1 µL of 20 mM AMP-PNP was added as well as 50 mM NaCl, 20 mM HEPES pH 7.4, 1 mM TCEP, 10% glycerol, 3 mM MgCl_2_, 0.025 mM ZnCl_2_ following incubation for 20 min, on ice, in a final volume of 20 µL. UV_254nm_ light crosslinking was performed using an in-house built crosslinking apparatus as previously described (38). Samples were irradiated for 10 min on ice. Mechlorethamine crosslinking was performed at a final concentration of 1 mM mechlorethamine (Sigma; 122564) for 30 min, 37°C, 300 rpm, followed by quenching with 50 mM Tris-HCl pH 7.5 and incubation for 5 min, room temperature, 300 rpm.UV- and mechlorethamine crosslinked samples were ethanol precipitated. After washing, pellets were dissolved in 4 M urea in 50 mM Tris-HCl pH 7.5. Samples were diluted to 1 M urea using 50 mM Tris/HCl pH 7.5. 1mM MgCl_2_, 250 U Pierce™ Universal nuclease for Cell Lysis (Thermo Scientific™, 88700), 100 U nuclease P1 (NEB, M0660S) and 1kU RNase T1 (Thermo Scientific™, EN0541) were added, following incubation for 3.75 hours, 37°C, 300 rpm. Trypsin (Promega; V5111) was added at a 1:20 enzyme-to-protein ratio following incubation overnight, 37°C, 300 rpm. Sample cleanup was performed using C18 stage tips (Harvard Apparatus; 74-4107) according to manufacturer’s instructions. Briefly, columns were equilibrated with 100% ACN; 50% [v/v] ACN, 0.1% [v/v] formic acid (FA); and three times 0.1% (v/v) FA. Samples were loaded twice and washed three times with 5% (v/v) ACN, 0.1% [v/v] FA. Elution was performed using 50% (v/v) ACN, 0.1% (v/v) FA and 80% (v/v) ACN, 0.1% [v/v] FA. Peptides were dried in a speed vac concentrator. Enrichment of crosslinked peptide-(oligo)nucleotides was performed as described (38) using Titansphere TiO_2_ 10 nm beads (GL Sciences; 5020-75010). After enrichment, samples were dried in a speed vac concentrator and subjected to LC-MS/MS measurements.

#### LC-MS/MS analysis

Enriched crosslinked peptides were injected onto a C18 PepMap100-trapping column (0.3 x 5 mm, 5 μm, Thermo Scientific™) connected to an in-house packed C18 analytical column (75 μm x 300 mm; Reprosil-Pur 120C18-AQ, 1.9 μm, Dr Maisch GmbH). Columns were equilibrated using 98% buffer A (0.1% [v/v] formic acid (FA)), 2% buffer B (80% (v/v) ACN, 0.1% (v/v) FA). Liquid chromatography was performed using an UltiMate-3000 RSLC nanosystem (Thermo Scientific™). Peptides were analyzed for 58 min using a linear gradient (5% to 45% buffer B (80% (v/v) ACN, 0.1% (v/v) FA in 44 min) followed by a 4.8 min washing step at 90% of buffer B. Eluting peptides were analyzed on an Orbitrap Exploris 480 instrument (Thermo Scientific™). The following MS settings were used: MS1 scan range, *m/z* 350–1600; MS1 resolution, 120,000 FWHM; AGC target MS1, 1E6; maximum injection time MS1, 60 ms; isolation window, *m/z* 1.4; top 30 most abundant precursors were selected for fragmentation; collision energy (HCD), 28% or 30% (for UV- and mechlorethamine-crosslinked samples, respectively);charge states, 2+ to 6+; dynamic exclusion, 7 s; MS2 resolution, 30,000, AGC target MS2, 1e5; maximum injection time MS2, 120 ms. The lock mass option (*m/z* 445.120025) was used for internal calibration.

#### Data Analysis

MS raw files were processed using the OpenMS framework for MS data analysis including NuXL tool (Urlaub group unpublished) and manual spectrum evaluation in TOPPView (39). For database search, the canonical protein sequence of MORC2 protein was used and default search settings with modifications, including variable modifications, oxidation (M), acetylation (N-term); enzyme, trypsin; maximum number of nucleotides attached, 2; and nucleotide adduct settings for UV or mechlorethamine crosslinking.

### Nucleosome preparation

A DNA consisting of 2 base pairs-Widom 601 sequence (145 base pairs)-2 base pairs was amplified by PCR using a 5’-/6-FAM forward primer (Sigma). The PCR products were pooled from four 96-well PCR plates (100 µL per well, 40 mL total volume) and isopropanol precipitated. DNA was purified using a 1 mL Resource Q column and eluted with a 22–34% NaCl gradient of TE Buffer (10 mM Tris pH 8.0, 2 M NaCl, 1 mM EDTA pH 8.0). Peak fractions were pooled, ethanol-precipitated, and resuspended in water. The 2-601-2 DNA sequence is as follows:5’-/6-FAM-ATA TCG ATG TAT ATA TCT GAC ACG TGC CTG GAG ACT AGG GAG TAA TCC CCT TGG CGG TTA AAA CGC GGG GGA CAG CGC GTA CGT GCG TTT AAG CGG TGC TAG AGC TGT CTA CGA CCA ATT GAG CGG CCT CGG CAC CGG GAT TCT GA T AT-3’

Nucleosomes were reconstituted essentially as described previously (40, 41). Histone octamer and DNA were mixed at a 1:1 molar ratio in RB-High Buffer (20 mM HEPES, pH 7.4, 2 M KCl, 1 mM EDTA, pH 8.0, and 1 mM DTT), transferred to Slide-A-Lyzer Mini Dialysis Units 20K MWCO, and gradient dialyzed against RB-Low Buffer (20 mM HEPES, pH 7.4, 30 mM KCl, 1 mM EDTA, pH 8.0, and 1 mM DTT) for 18 hours at 4 °C. The sample was further dialyzed against RB-Low Buffer for 2 hours at 4 °C. The sample was centrifuged for 1 min at 11,000 rpm to collect precipitate. Nucleosome concentration was quantified by absorbance at 280 nm. The molar extinction coefficient of the nucleosome was obtained by summing the molar extinction coefficients of the octamer and the DNA components at 280 nm.

### Fluorescence anisotropy

5’-/6-FAM labeled double-stranded DNA oligos were obtained from IDT, 5’-/6-FAM labeled single-stranded DNA and single-stranded RNA were obtained from Sigma Aldrich and dissolved in water to 100 µM. Sequences of nucleic acid substrates used are detailed in the table below.

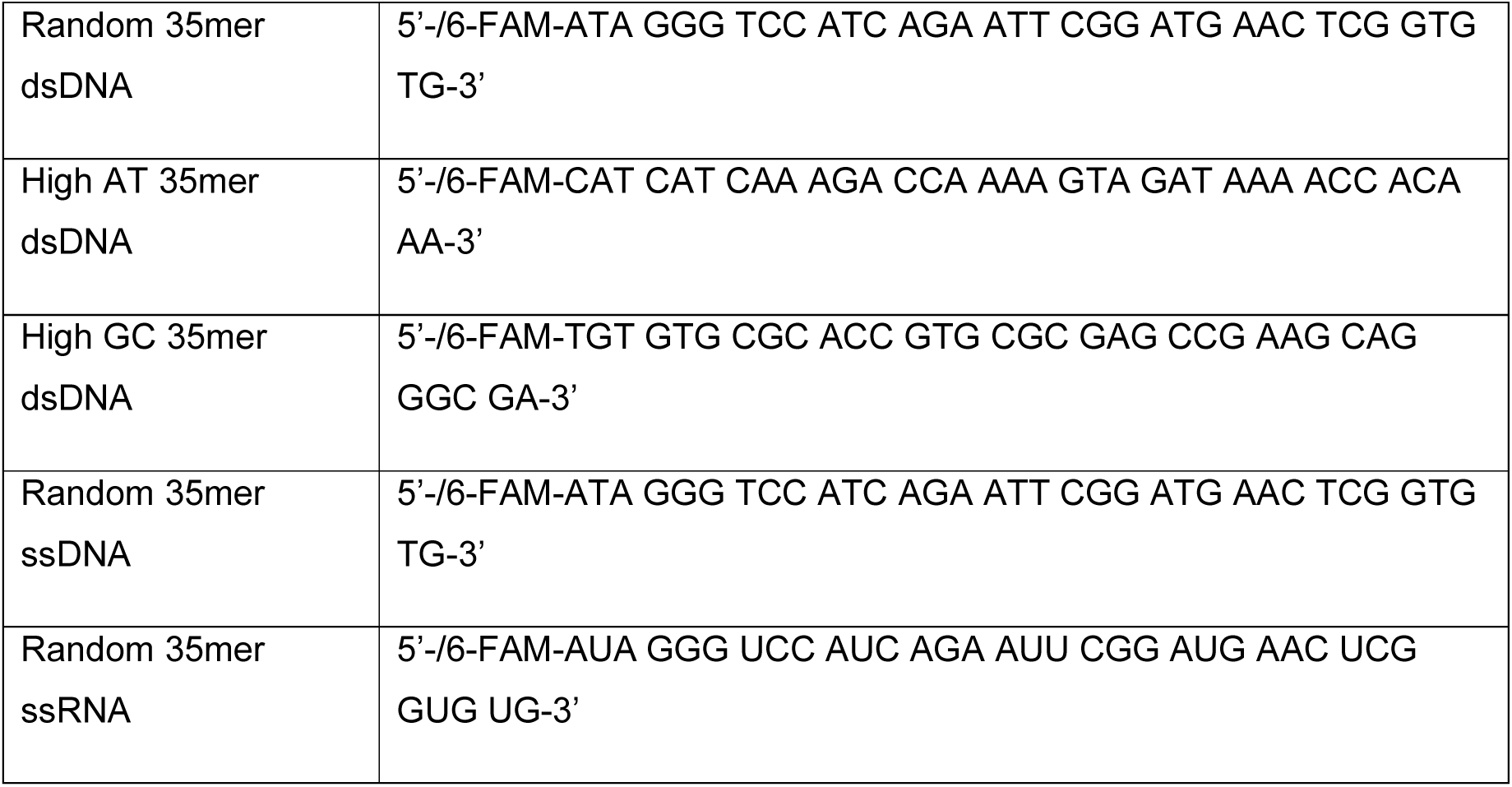
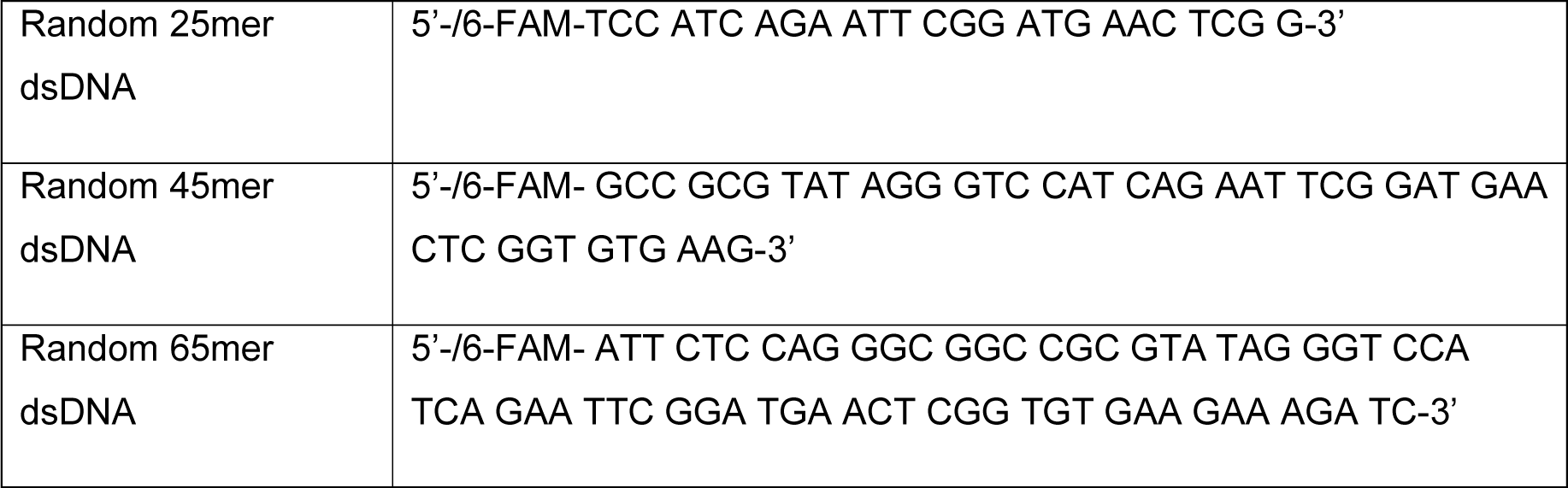

MORC2 protein was diluted in half in Half Dilution buffer (20 mM Na•HEPES pH 7.4, 10% (v/v) glycerol, and 1 mM TCEP) and then serially diluted in half log steps in Protein Dilution buffer (250 mM NaCl, 20 mM Na•HEPES pH 7.4, 10% (v/v) glycerol, and 1 mM TCEP). Nucleic acid (5 µL, 1 nM final concentration) was mixed with MORC2 dilutions (5 µL, 1-0.001 µM final concentrations) on ice and incubated for 10 minutes. The assay was brought to a final volume of 25 µL and incubated in the dark at room temperature for 20 minutes (final conditions: 50 mM NaCl, 3 mM MgCl_2_, 20 mM Na-HEPES pH 7.4, 1 mM TCEP, 10% (v/v) glycerol, 0.025 mM ZnCl_2_ and 50 µg/ml BSA). 18 µL of each solution was transferred to a Greiner 384 Flat Black Bottom plate.

Fluorescence anisotropy was measured at room temperature with a Tecan SPARK plate reader with an excitation wavelength of 470 nm (±5 nm), an emission wavelength of 518 nm (±20 nm), a gain of 150, and a Z-height of 26050 µm. Experimental measurements with protein were normalized to buffer with nucleic acids alone. All experiments were done in triplicate. Binding curves were fit with a single site quadratic binding equation:

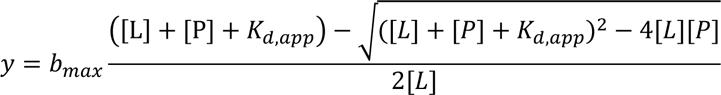

where B_max_ is the maximum specific binding, L is the concentration of nucleic acid, P is the concentration of MORC2, and K_d,app_ is the apparent dissociation constant. Curve fitting was performed in GraphPad Prism (10.0.3).

### Malachite Green ATPase assay

MORC2 ATP hydrolysis activity was measured using a malachite green assay. MORC2 protein was initially diluted in half in Half Dilution buffer (20 mM Na•HEPES pH 7.4, 10% (v/v) glycerol, and 1 mM TCEP) and then diluted to 5 µM in Protein Dilution buffer (250 mM NaCl, 20 mM Na•HEPES pH 7.4, 10% (v/v) glycerol, and 1 mM TCEP). MORC2 (10 µL, 1 µM final concentration) was mixed with water or 35 base pair random dsDNA substrate (10 µL, 2 µM final concentration or as indicated) on ice for 10 minutes. The assay was brought to a volume of 45 µL and incubated at 37°C for 5 minutes (final conditions: 50 mM NaCl, 3 mM MgCl_2_, 20 mM Na-HEPES pH 7.4, 1 mM TCEP, 10% (v/v) glycerol, 0.025 mM ZnCl_2_, 2 mM phosphoenol pyruvate, 4 U Pyruvate Kinase/ Lactate Dehydrogenase, and 50 µg/mL BSA). Ultrapure ATP (Jena Bioscience) was added to the reaction (5 µL, 1 mM final concentration or as indicated). After incubating the reactions for 45 minutes at 37°C, reactions were quenched with EDTA pH 8.0 (20 mM final concentration) and moved to ice. For ATPase measurements where ATP was titrated, reactions were prepared as described above except with 10 mM MgCl_2_ final concentration to provide enough Mg^2+^ ions to coordinate the excess ATP in the reaction. For each experiment, a titration of sodium phosphate from 5-500 µM in the final reaction conditions was made to generate a standard curve.

Sulfuric acid (320 mM final concentration) was added to precipitate protein on ice for 10 minutes. The malachite green detection mixture was prepared on ice (1 mL 0.12% (w/v) malachite green, 330 µL 14% (w/v) ammonium molybdate, 20 µL 11% (v/v) Tween-20). Quenched and precipitated samples were centrifuged at 15,000 rpm for 1 minute. 50 µL of samples were transferred to a Greiner 384 Flat Clear Bottom plate. 25 µL of malachite green detection mixture was added for a two-minute incubation in the dark and then 7.5 µL 34% (w/v) sodium citrate was added for an additional 28-minute incubation in the dark. Absorbance at 620 nm was measured in a Tecan SPARK plate reader at room temperature. Experimental measurements with protein were normalized to a buffer control with substrate alone. All experiments were done in triplicate. The standard curve was used to convert absorbance values to the concentration of free phosphate. The concentration of free phosphate was assumed to equal the amount of ATP hydrolyzed by MORC2. ATP hydrolysis rates are reported as µM ATP hydrolyzed per µM MORC2 protein per minute. The ATP titration data are fit to the Michaelis-Menten model:

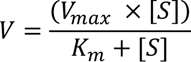

where V is the reaction rate, V_max_ is the maximum rate, [S] is the total starting substrate concentration, and K_m_ is the Michaelis-Menten constant. Curve fitting was performed in GraphPad Prism (10.0.3). The DNA titration data were fit to an inhibition curve:

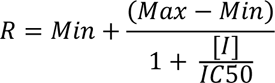

where R is the rate, Min is the minimum rate, Max is the maximum rate, [I] is the inhibitor concentration, and IC50 is the inhibition constant. Curve fitting was performed in GraphPad Prism (10.0.3).

### Size-exclusion chromatography (SEC)- coupled to multi-angle light scattering (MALS)

SEC-MALS analysis of MORC2 constructs was performed using 100 µL of protein in final conditions: 3 mM MgCl_2_, 20 mM Na-HEPES pH 7.4, 0.5 mM TCEP, 1.5% (v/v) glycerol, and 0.025 mM ZnCl_2_. For full-length MORC2, 2 mg/mL of MORC2 was prepared in 150 mM NaCl with or without the addition of 1 mM AMP-PNP. For 1-603 MORC2, 1 mg/mL of 1-603 MORC2 was prepared in 50 mM NaCl in the presence or absence of 1 mM AMP-PNP. Samples were analyzed on an analytical 030 column (Wyatt) run at a flow rate of 0.5 mL/min. 1.5 mg/mL of coiled coil 3 protein was prepared in 150 mM NaCl and was analyzed on an analytical 010 column (Wyatt). Light scattering analysis was performed in ASTRA (Wyatt), using band broadening parameters obtained from a BSA standard in identical running conditions. MALS data were used to fit the average molar mass across the peak of interest, reported to the nearest kDa.

### Plasmid capture assay

Circular plasmid substrates, pUC19 and pBlueScript, were prepared by maxiprep following the manufacturer’s instructions (Qiagen). Linear pUC19 substrate was prepared by incubating 50 µg of pUC19 with EcoRI for 6 hours at 37°C. Linear pUC19 was purified by phenol chloroform extraction and precipitated with isopropanol.

MBP-tagged MORC2 was diluted in half in Half Dilution buffer (20 mM Na•HEPES pH 7.4, 10% (v/v) glycerol, and 1 mM TCEP) and then further diluted in Protein Dilution buffer (250 mM NaCl, 20 mM Na•HEPES pH 7.4, 10% (v/v) glycerol, and 1 mM TCEP). Diluted MORC2 (600 nM final, monomer concentration) was combined with circular or linear substrate (100 nM final) and incubated at room temperature for 20 minutes. The assay was brought to 15 µL final volume in the presence or absence of 1 mM AMP-PNP and incubated at room temperature for 30 minutes (final conditions: 50 mM NaCl, 3 mM MgCl_2_, 20 mM Na-HEPES pH 7.4, 1 mM TCEP, 10% (v/v) glycerol, 0.025 mM ZnCl_2_ and 50 µg/ml BSA). 50 µL amylose resin was washed with water and equilibrated in Reaction Buffer (50 mM NaCl, 3 mM MgCl_2_, 20 mM Na-HEPES pH 7.4, 1 mM TCEP, 10% (v/v) glycerol, 0.025 mM ZnCl_2_ and 50 µg/ml BSA). 1 µL of the reaction was saved as an input sample and the remaining 14 µL of the reaction was applied to the equilibrated amylose resin. The reaction mixture was incubated with the amylose resin for 30 minutes at room temperature shaking at 800 rpm. The mixture was washed twice with 250 µL of either Reaction Buffer or a High Salt Buffer (400 mM NaCl, 3 mM MgCl_2_, 20 mM Na-HEPES pH 7.4, 1 mM TCEP, 10% (v/v) glycerol, and 0.025 mM ZnCl_2_). Samples with AMP-PNP were washed with buffer supplemented with 1 mM AMP-PNP. Protein was eluted from the resin with 20 µL Maltose Elution buffer (50 mM NaCl, 3 mM MgCl_2_, 20 mM Na-HEPES pH 7.4, 1 mM TCEP, 10% (v/v) glycerol, 0.025 mM ZnCl_2_, 50 µg/ml BSA, and 100 mM maltose). Reactions were quenched with EDTA (10 mM final). Input samples were diluted to 10 µL with Reaction buffer. Protein in all samples and input samples was digested by proteinase K. All samples were supplemented with 10% sucrose and visualized on 1% (w/v) 1x TAE (50 mM Tris-HCl, pH 7.9, 1 mM EDTA, pH 8.0, 40mM sodium acetate) agarose gels with 1:10,000 SYBR Safe stain (Apex) run at 100 V for 50 minutes. Gels were imaged on a BioRad Gel Dock imager with a 1.5 second exposure.

Gel images were quantified in ImageJ (1.52A). A 60x40 pixel box was drawn around the input or retained DNA band for each lane and the integrated density inside the box was measured. For each lane, a 60x40 pixel box was drawn below the DNA band and the integrated density inside the box was measured as the background. The reported values are the difference between the integrated density measurements of the DNA and the background for each lane.

### Dual DNA capture assay

DNA bead stocks were prepared essentially as described before (42). pUC19 DNA was amplified by PCR with the following primers: 5’-biotin-CGG TGA AAA CCT CTG ACA CAT G-3’ and 5’-biotin-TCA TCA CCG AAA CGC GC-3’. The desired product was separated from PCR byproducts and primers on a 1 mL resource Q column. Samples were loaded on the column in TE Zero buffer (10 mM Tris-HCl, pH 8.0, 1 mM EDTA, pH 8.0), washed with 28% TE High buffer (10 mM Tris-HCl, pH 8.0, 1 mM EDTA, pH 8.0, 2M NaCl), and eluted by a gradient from 28% - 34% TE High buffer. Fractions containing the desired product were pooled and DNA was isopropanol precipitated. Dynabeads M-280 Streptavidin beads (Invitrogen) were washed three times in DNA Binding buffer (10 mM HEPES, pH 7.4, 2M NaCl, 1 mM EDTA, pH 8.0). 2.5 µg of DNA per 100 µL of beads were added and the mixture was diluted in half with water. Samples were incubated overnight at room temperature with gentle rotation. Beads were washed three times in Reaction Buffer and stored at 4°C.

MORC2 was diluted in half with Half Dilution buffer (20 mM Na•HEPES pH 7.4, 10% (v/v) glycerol, and 1 mM TCEP) and then further diluted in Protein Dilution buffer (250 mM NaCl, 20 mM Na•HEPES pH 7.4, 10% (v/v) glycerol, and 1 mM TCEP). Diluted MORC2 (600 nM final) was combined with 20 µL DNA beads and incubated at room temperature for 20 minutes with shaking at 800 rpm in the presence or absence of 1 mM AMP-PNP. pBlueScript (200 nM final) was added and the mixture was incubated at room temperature for 30 minutes, shaking. Samples were washed twice with 250 µL of either Reaction Buffer or High Salt Buffer (400 mM NaCl, 3 mM MgCl_2_, 20 mM Na-HEPES pH 7.4, 1 mM TCEP, 10% (v/v) glycerol, and 0.025 mM ZnCl_2_). Samples with AMP-PNP were washed with buffer supplemented with 1 mM AMP-PNP. Beads were resuspended in 1X CutSmart Buffer (New England Biolabs), and DNA was digested with 20 U ScaI-HF and 20 U Sbf-HF for 1 hour at 37°C. Supernatants were removed and protein was digested with proteinase K for 1 hour at 37°C. All samples were supplemented with 10% sucrose and visualized on 1% (w/v) 1x TAE (50 mM Tris-HCl, pH 7.9, 1 mM EDTA, pH 8.0, 40mM sodium acetate) agarose gels with 1:10,000 SYBR Safe stain (Apex) run at 100 V for 50 minutes. Gels were imaged on a BioRad Gel Dock imager with a 1.5 second exposure.

Gel images were quantified in ImageJ (1.52A). A 60x30 pixel box was drawn around the input bead DNA band and the retained second plasmid DNA band for each lane and the integrated density inside the box was measured. For each lane, a 60x30 pixel box was drawn below the input bead DNA band and the integrated density inside the box was measured as the background. The reported values are the ratio of the difference between the integrated density measurements of the retained second plasmid DNA and the background for each lane and the difference between the integrated density measurements of the input bead DNA and the background for each lane.

### Circular dichroism

MORC2 protein samples were dialyzed overnight at 4°C with stirring against CD Dialysis Buffer (100 mM NaCl, 10 mM HEPES, pH 7.4). Samples were retrieved and diluted to 0.1 mg/mL in CD Dialysis Buffer. CD spectra were taken at 25°C with a JASCO model J-1500 Circular Dichroism Spectrophotometer in a 1-mm pathlength quartz cuvette. Data were obtained from 250 to 200 nm, at 0.5 nm intervals. Each point is averaged over 4 sec, and each read was performed three times. Reported values are the difference between the protein sample and the buffer alone control.

### NLS Stradmus analysis

NLS Stradmus tool for nuclear localization signal prediction was described previously (43). For MORC2, a 2-state static Hidden Markov Model was used, and sequences were selected above a 0.4 posterior threshold cutoff.

### Multiple Sequence Alignment

Primary sequences of MORC2 proteins from various organisms were obtained from Uniprot (IDs: Q9Y6X9, K7D393, F1RPD9, M3W1K4, A0A8C0I0T3, Q69ZX6, D4A2C4, A0A1L8HZT1, A0A670ZP14, Q68EG7). Sequences were aligned using MAFFT (version 7) and visualized in Jalview (2.11.3.0).

### AlphaFold Multimer

The AlphaFold Multimer dimer structure of full-length MORC2 was predicted using the COSMIC2 server (44). Structure was visualized in PyMOL (2.5.2).

### Cell culture conditions

MORC2 knockout HeLa cells were generated by Ubigene Biosciences using a CRISPR gene editing strategy. Briefly, HeLa cells were electroporated with Cas9 protein and gRNAs flanking exon 5 of the MORC2 gene, followed by single clone selection. Sequences of the gRNAs are detailed below with the PAM sequence italicized.

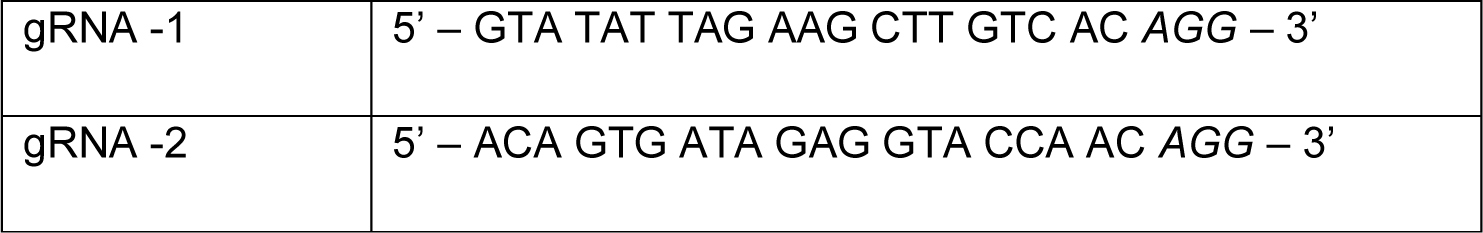

To genotype MORC2 knockouts, cells were lysed in DirectPCR lysis reagent (Viagen, 301-C) with proteinase K (0.2 µg/mL) overnight at 55°C then proteinase K was inactivated at 85°C for 90 minutes. Clones were screened by PCR amplification of a region of the MORC2 gene around exon 5 such that removal of exon 5 would result in a shorter PCR product. The resulting PCR product was also validated by Sanger sequencing (Quintara). Sequences of the primers used for PCR amplification are detailed below.

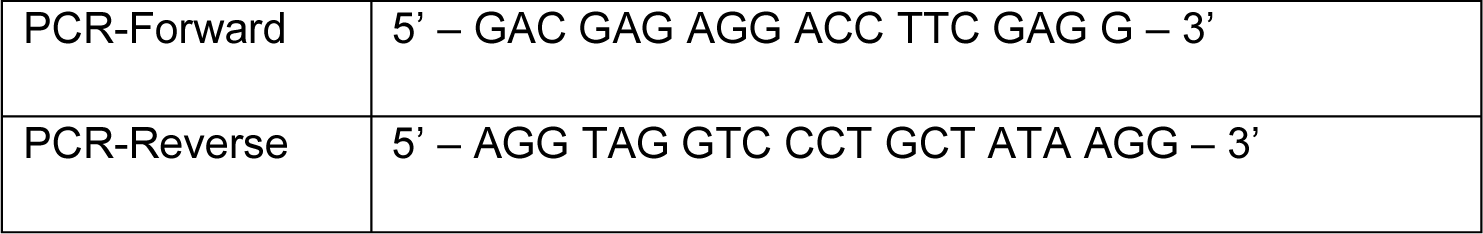

HeLa cell lines (a gift from the lab of Iain Cheeseman) and MORC2 knockout HeLa cell lines were cultured in DMEM supplemented with 10% tetracycline-free fetal bovine serum (FBS), 2 mM L-glutamine, and 100 U/mL penicillin/streptomycin at 37°C with 5% CO_2_. The piggyBac transposon system (45) was used to introduce EGFP-tagged MORC2 constructs to wildtype or MORC2 knockout HeLa cells. Cells were plated in 6-well plates. To generate stable tetracycline inducible cell lines, 1 µg donor plasmid containing the tetracycline inducible transgene was co-transfected with 400 ng piggyBac transposase plasmid (HP137, a gift from the lab of Rudolf Jaenisch) using lipofectamine 2000. Cells were grown at 37°C for 1 day, then the media was exchanged. 48 hours post transfection, cells were moved into a 10 cm^2^ plate and selected for puromycin resistance using puromycin 0.35 µg/ mL for 5 days. Cell lines were frozen at approximately 1 million cells/mL in DMEM supplemented with 25% FBS and 10% DMSO and stored in liquid nitrogen.

### Protein extraction and Western blots

Pellets of approximately 5 million HeLa cells were thawed on ice and resuspended in lysis buffer (2X TBS, 1% Triton X-100, 1X HALT protease and phosphatase inhibitor cocktail). Protein concentrations were quantified using BCA Protein Assay Kit (Thermo Scientific) following the manufacturer’s protocol. 50 µg of total proteins were loaded into each lane of a 4-12% Bis-Tris gradient gel (NuPAGE), and transferred to 0.45 µm PVDF membranes (GE Healthcare). Membranes were blocked with 5% BSA in TBST (Tris buffered saline pH 8.0, 0.1% Tween 20, Thermo Scientific) for 1 hour at room temperature and incubated with primary antibody overnight at 4°C. Membranes were washed with TBST and incubated with secondary HRP conjugate antibodies for 1 hour at room temperature, followed by washing with TBST and development using a chemiluminescence substrate (Thermo Fisher #34580). Membranes were exposed using Bio-rad Chemidoc^TM^. Primary antibodies used: β-ACT (Sigma-Aldrich #A5316, 1:10,000 dilution in 5% BSA in TBST), MORC2 (Bethyl Laboratories #A300-149A, 1:1,000 dilution in 5% BSA in TBST). Secondary antibodies used: anti-mouse IgG HRP-linked (Invitrogen #62-6520, 1:10,000 dilution in 5% BSA in TBST), anti-rabbit IgG HRP-linked (CST #7074, 1:10,000 dilution in 5% BSA in TBST).

### Fluorescence microscopy

Cells were plated in 12-well glass-bottom plates and treated with 1 µg/ mL doxycycline for 24 hours. Medium was replaced with CO_2_-independent medium with 0.1 µg/ mL Hoechst to visualize DNA and incubated on the stage for 30 minutes before imaging. Cells were maintained at 37°C using a heated stage. Images were taken on a Deltavision Ultra microscope (Cytiva) using a 60x/1.42NA objective. 8 μm images were taken with z sections of 0.2 μm. All images shown are deconvolved and maximally projected (2 μm) using Fiji/ImageJ. For images where intensities were compared between samples, the images were taken with identical exposure settings, intensity, and contrast were scaled equivalently (Figure 5C). For images where the localization of the protein was being assessed, the images were not scaled equivalently (Figure 5A, Supplementary Figure S7B). Morphological markers such as DNA were not scaled equivalently.

### Quantification of live-cell images

The relative mitotic chromosome-cytoplasmic ratios of EGFP-MORC2 constructs were quantified with a custom automated CellProfiler (v4.2.6) (46) pipeline. The pipeline segmented mitotic chromosomes and cytoplasm based on Hoescht and EGFP signal and measures mean intensity. Chromosomes were detected with the options: min/max diameter of 75 - 500, adaptive otsu thresholding method with two classes. The size of the adaptive window is 50 without log transformation before thresholding. To identify the cell boundary, we used the EGFP signal with the watershed – image method and a global minimum cross-entropy thresholding method. The mean EGFP intensity was measured within the chromosome and in the cytoplasm (excluding the chromosome). Image quantification was performed on raw, non-deconvolved, maximally projected images. The chromosome/cytoplasm ratio was normalized to wildtype EGFP-MORC2.

### RNA extraction and sequencing

Cells were plated in 6-well plates and treated with 1 µg/ mL doxycycline for 48 hours. Cells were washed once with PBS, resuspended 400 µL of TRI Reagent (Invitrogen, AM9738), and stored at -80°C. 120 µL of chloroform was added to the TRI reagent, the mixture was vortexed then separated by centrifugation at 21000x g, 4°C for 15 minutes. The aqueous layer containing the nucleic acids was collected. Nucleic acids were precipitated with 300 mM NaCl and 30 µg GlycoBlue (AM9516) by the addition of equal volumes of isopropanol and incubation at -20°C overnight. Precipitated nucleic acids were collected by centrifugation at 21000x g, 4°C for 15 minutes. The supernatant was removed, and the nucleic acid pellet was washed with 70% ethanol and dissolved in RNase-free water. Nucleic acid concentration was quantified using the Qubit RNA Broad Range kit (Q10210).

To generate spike-in mRNAs for absolute RNA quantification in RNA-seq, plasmids containing T7 promoter – Nano or Firefly Luciferase were amplified by PCR. The template for PCR was digested with DpnI then purified using EconoSpin TM All-in-1 Mini Spin Columns (Epoch Life Science, 1920-250) and eluted in RNase free water. 500 ng of the PCR product was used in an in vitro transcription reaction using the HiScribe T7 High Yield RNA Synthesis Kit (NEB, E2040S). The DNA template was digested through addition of DNase I after in vitro transcription. Free nucleotides were removed using a P30 spin column (Bio Rad, 7326251) then the RNA was purified by phenol chloroform extraction and isopropanol precipitation. The RNA was capped using the Vaccinia Capping System (NEB, M2080S), desalted on P30 columns, phenol chloroform extracted, and precipitated overnight at -20°C with isopropanol. The capped RNA was polyadenylated using *E. coli* polyA polymerase (NEB, M0276L), desalted on P30 columns, phenol chloroform extracted, and precipitated overnight at -20°C with isopropanol. The purified capped and polyadenylated mRNA was resuspended in RNase free water, the Nano and Firefly luciferase was mixed at a ratio of 1:2, aliquoted, flash frozen in liquid nitrogen, and stored at -80C.

For RNA-sequencing, 2 µg of total RNA was supplemented with 5 pg in vitro transcribed Nano and Firefly luciferase mRNA. RNA integrity was assessed using Agilent’s Fragment Analyzer prior to library preparation. Sequencing libraries were prepped using the NEBNext Ultra II RNA Library Prep Kit and sequenced using the Element AVITI sequencer with 75x75 bp paired end runs.

### Analysis of RNA-sequencing data

A custom genome and annotation containing nano luciferase mRNA, firefly luciferase mRNA, transposable elements (hg38, repeatmasker), and human canonical protein coding mRNAs (hg38, gencode v35) was used for mapping. The custom genome was indexed using STAR (v2.7.1a) (47) with the option --sjdbOverhang 74. As suggested by Tetranscripts (48), reads were aligned to the hg38 reference genome using STAR with options –outFilterMultimapNmax 100 --winAnchorMultimapNmax 200 --outFilterType BySJout --outSAMattributes All -- outSAMtype BAM SortedByCoordinate. Mapped .bam files were indexed and reads were quantified using Tetranscripts (v2.2.3) (48) with the following options --sortByPos --format BAM --mode multi --stranded reverse --minread 25. For non-spike normalized data (Supplementary Figure S9C), the DEseq2 output from the TEtranscript package was used, where the default DEseq2 median of ratios method for normalization was performed.

For spike-in mRNA quantification, the mapped reads were quantified using htseq-count (v1.99.2) (49) with the options -r pos --stranded=reverse -f bam -t exon. The sample sizeFactor was calculated by summing counts of nano and firefly luciferase mRNA then normalizing the summed spike mRNA counts to control NLS-GFP replicate 1. These sizeFactors along with raw count outputs from TEtranscripts were used for DEseq2 (50) to calculate adjusted p-values and fold changes.

For PCA analysis (Supplementary Figure S9A), the spike normalized DEseq2 data was rlog transformed and PCA plots were generated using plotPCA with default settings from the DEseq2 package. For quantification of MORC2 mRNA expression from RNA sequencing data (Supplementary Figure S9B), the raw counts from TEtranscripts were normalized using DEseq2 median of ratios.

## Results

### MORC2 binds DNA and phosphorylation reduces its affinity for DNA

To characterize the interaction of full-length human MORC2 with DNA, we recombinantly overexpressed human MORC2 using a baculovirus expression system and purified it to homogeneity (**Figure 1B, Supplementary Table S1, Methods**). Phosphorylation site analysis by mass spectrometry of the purified protein indicates that MORC2 is heavily phosphorylated during its expression in insect cells (**Methods**). We next assessed how phosphorylated and dephosphorylated MORC2 associate with DNA using fluorescence anisotropy (FA) with a 5’ FAM labelled 35mer double stranded DNA (dsDNA) oligo. Dephosphorylated MORC2 binds DNA with ∼10 fold higher affinity than MORC2 preparations where phosphorylations remain intact (phosphorylated: K_d,app_ 285 ± 83 nM, dephosphorylated: K_d,app_ 17 ± 4 nM) (**Figure 1C**).

We identified seventeen phosphorylation sites that are known MORC2 phosphorylation sites in human cell lines (S615, T650, S696, S703, S705, S711, T717, T723, S725, S730, T733, S735, S739, S743, S773, S777, S779) (**Figure 1A**) (51). Interestingly, three phosphorylation sites (S725, S730, and S773) were identified as residues under positive selection during primate evolution (31). We mutated the seventeen identified serine and threonine phosphorylation sites to aspartate and glutamate, respectively, to mimic the expected charge in a fully phosphorylated state (phosphomimetic). We also prepared a construct where the phosphorylation sites were substituted to alanine to prevent phosphorylation (phosphodead). Both mutants appear to be well-folded and hydrolyze ATP (**Supplementary Figure S1B, Supplementary Figure S4A**). The phosphomimetic mutant has a greatly reduced affinity for DNA (K_d,app_ 512 ± 147 nM) whereas alanine substitutions marginally affect DNA association (K_d,app_ 25 ± 3 nM) (**Figure 1C**). Together, these results suggest that phosphorylation of MORC2 in the region between residues 603-790 impairs DNA association. For the remainder of this work, we used dephosphorylated MORC2 preparations.

In addition to DNA, MORC3 has been shown to bind single stranded RNA and nucleosomal substrates (19, 52). There is also evidence that MORC3 associates with specific DNA sequences (8, 53) and that *C. elegans* MORC-1 preferentially binds longer DNA substrates (>1000 bp) (15). We thus tested whether full-length MORC2 has a binding preference for dsDNA, single stranded DNA (ssDNA), single stranded RNA (ssRNA), a nucleosomal substrate, dsDNA substrates containing different GC contents, and dsDNAs of different lengths. Nucleosomes were reconstituted *in vitro* by combining purified histone octamer and a 149 bp dsDNA with a Widom 601 nucleosome positioning sequence (54). MORC2 most readily associates with double stranded substrates, with a slight preference for the 149 bp Widom 601 dsDNA than the reconstituted nucleosome (149 bp Widom 601 dsDNA: K_d,app_ 5 nM or less, nucleosome: K_d,app_ 16 ± 2 nM,) (**Figure 1E**). The MORC2 affinity for ssDNA or ssRNA is about 10-fold less than its affinity for dsDNA (ssDNA: K_d,app_ 51 ± 9 nM, ssRNA: K_d,app_ 49 ± 6 nM). We also observe that GC content of the DNA substrate does not substantially affect MORC2 DNA association, and MORC2 appears to preferentially associate with longer DNAs (**Figure 1F, Supplementary Figure S4C, Supplementary Table S2**). These results suggest that full-length MORC2 preferentially binds dsDNA with no apparent sequence preference.

### A C-terminal region of MORC2 strongly associates with DNA

To identify regions of MORC2 that associate with DNA, we performed mechlorethamine and UV-induced DNA-protein cross-linking with dephosphorylated MORC2 and a 65mer dsDNA oligo (**Methods**). Crosslinks were detected by mass spectrometry. Mechlorethamine primarily crosslinks purines with amino acids that are 7-8 Å away whereas UV light induces zero-length crosslinks between amino acids adjacent to thymidines, primarily. We identified regions of MORC2 in both the N- and C-terminus that form multiple crosslinks to DNA with both crosslinking approaches (**Figure 1A**, **Supplementary Table S3**). Notably, a few of the N-terminal crosslinks surround a previously identified region in coiled coil 1 (residues 326-333) which has been shown to associate with DNA (18). To test the relative DNA binding contributions of the N- and C-terminus, we purified an N-terminal fragment of MORC2 (residues 1-603). The N-terminal MORC2 fragment binds DNA weakly (K_d,app_ 446 ± 119 nM) as reported previously (**Figure 1D**) (18).

Four of the C-terminal crosslinking sites abut a positively charged region bearing twelve highly conserved lysine and arginine residues (K704, R707, K713, K716, K721, K722, R754, R755, K756, R758, K760, R761), suggesting that these residues could mediate DNA binding (**Figure 1A, Supplementary Figure S2, Supplementary Figure S3**). To test this idea, we mutated these residues to either aspartate or alanine and purified the resulting full-length mutants. Both the aspartate and alanine mutant bind DNA with a greatly reduced affinity compared to the wildtype protein (aspartate mutant: K_d,app_ 380± 120 nM, alanine mutant: K_d,app_ 350± 60 nM) (**Figure 1D, Supplementary Figure S4B**). We next mutated subsets of the positively charged residues to aspartate (N-terminal subset (subset N): K704, R707, K713, K716, K721, K722 and C-terminal subset (subset C): R754, R755, K756, R758, K760, R761). Partial removal of the positively charged residues results in an intermediate reduction in MORC2 DNA association (subset N: K_d,app_ 90 ± 20 nM, subset C: K_d,app_ 60 ± 20 nM) (**Supplementary Figure S4B**). Together, these results suggest that a region corresponding to MORC2 704-761 is required for strong DNA association. Notably, this region overlaps with the identified phosphorylation sites which reduce DNA binding.

### MORC2 ATP hydrolysis activity is reduced in the presence of DNA

The rate of ATP hydrolysis by DNA transacting ATPases can be positively or negatively influenced by their association with DNA (19, 20, 55–58). We thus tested whether DNA affects MORC2 ATP hydrolysis activity. To do this, we utilized a malachite green endpoint assay that detects the release of inorganic phosphate. The MORC2 1-603 construct hydrolyzes ATP with a rate of 0.09 ± 0.02 min^-1^, which is in agreement with prior work (**Figure 2A**) (18). Full-length MORC2, in the absence of DNA, hydrolyzes ATP four times faster than MORC2 1-603 (0.39 µM ± 0.05 min^-1^). Mutations which should either alter ATP binding (N39A) or hydrolysis (E35A) result in almost no detectable ATPase activity (**Figure 2A**) (18, 59). We next included our 35mer dsDNA oligo at a 4:1 molar ratio DNA:MORC2 dimer in the assay and observed an almost four-fold reduction in MORC2 ATPase rate (0.11 ± 0.06 min^-1^). DNA inhibits MORC2’s ATPase rate in a dose dependent manner (DNA IC_50_ 0.14 ± 0.05 µM, [MORC2 dimer] = 0.5 µM) (**Figure 2B**). In contrast, DNA does not appear to influence the rate of ATP hydrolysis by MORC2 1-603 (0.12 ± 0.01 min^-1^) or the DNA binding deficient aspartate MORC2 mutant (without DNA 0.20 ± 0.04 min^-1^, with DNA 0.19 ± 0.02 min^-1^) (**Figure 2A**).

**Figure 2.**
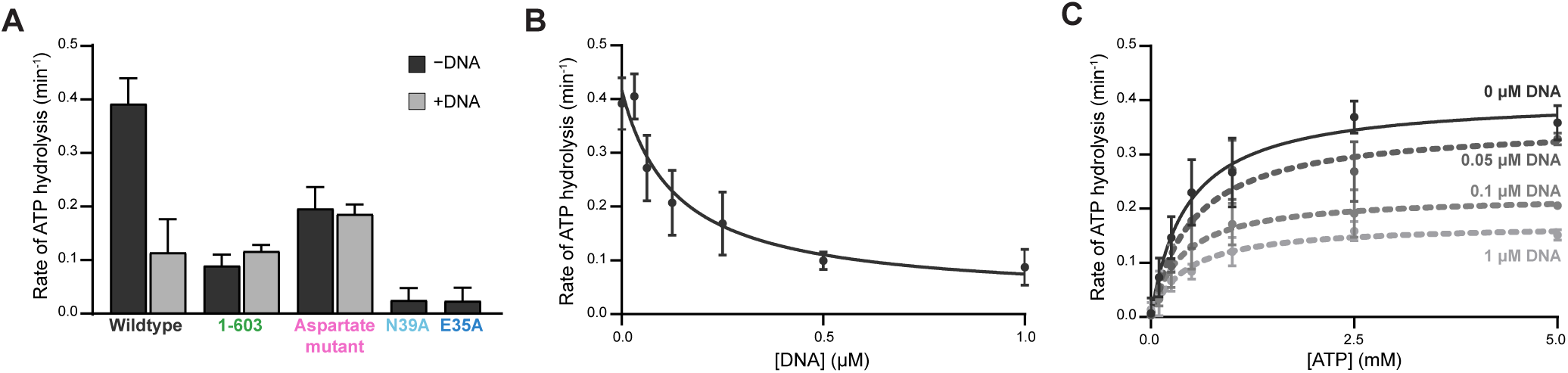
DNA binding by MORC2 reduces ATPase activity. **A.** Assessment of MORC2 ATPase activity. Wildtype, 1-603, DNA binding deficient aspartate mutant, and ATP hydrolysis deficient mutant E35A, and ATP binding deficient mutant N39A MORC2 (1 µM) were incubated with 1 mM ATP for 45 minutes at 37°C either in the presence or absence of 2 µM of a 35 base pair DNA. Inorganic phosphate released was quantified by malachite green (**Methods**). Error bars correspond to the standard deviation between three replicate experiments. **B.** MORC2 ATPase activity is reduced in the presence of DNA. 1 µM MORC2 was incubated with 1 mM ATP for 45 minutes at 37°C with a titration of a 35 base pair duplex DNA. Inorganic phosphate released was quantified by malachite green (**Methods**). Data were fit to an inhibition curve. Error bars correspond to the standard deviation between three replicate experiments. **C.** Michaelis-Menten analysis of MORC2 ATPase activity in the presence of DNA. 1 µM MORC2 was incubated with an ATP titration for 45 minutes at 37°C in the presence of various concentrations of 35mer DNA (0-1 µM). Inorganic phosphate released was quantified by malachite green (**Methods**). Data were fit to a Michaelis-Menten model of enzyme kinetics. Error bars correspond to the standard deviation between three replicate experiments.

We next tested how DNA affects the kinetics of ATP hydrolysis by MORC2 by measuring K_m, app ATP_ and V_max_ of the enzyme with varying concentrations of DNA. As DNA concentration increases, K_m, app ATP_ remains constant whereas V_max_ decreases (**Figure 2C, Supplementary Table S4**). Our K_m, app ATP_ for full-length MORC2 corresponds to the previously reported K_m, app ATP_ for MORC2 1-603 (18). Together, these results suggest that DNA non-competitively inhibits MORC2 ATPase activity, and DNA binding may induce a conformational change in MORC2 that results in reduced ATP hydrolysis.

Finally, since MORC2 ATP hydrolysis is reduced in the presence of DNA, we tested whether ATP or ATP analogs affect DNA association with MORC2. All tested nucleotides result in a ∼5-50-fold reduction in DNA binding affinity (**Supplementary Figure S4D, Supplementary Table S2**). Intriguingly, ATPγS leads to the largest reduction in DNA binding affinity. It has been shown that the MORC3 GHKL dimer interface is most strongly stabilized by ATPγS (60). One possible explanation for these observations is that dimerization of the GHKL domain in the presence of ATP or ATP analogs limits MORC2 association with DNA.

### MORC2 contains N- and C-terminal dimerization interfaces

GHL-type ATPases contain a second, C-terminal dimerization interface (61). To determine the oligomeric state of MORC2 1-603 and full-length MORC2, we utilized size exclusion chromatography coupled to multi-angle light scattering (SEC-MALS). We observe that the MORC2 1-603 only dimerizes in the presence of ATP or ATP analogs, as previously observed (18). In contrast, full-length MORC2 forms a dimer in both the presence and absence of the ATP analog AMP-PNP. These results suggest the presence of a second, C-terminal dimerization interface (**Figure 3A**). An AlphaFold multimer prediction of the full-length, dimer structure indicates that the C-terminal coiled coil 3 domain could make an additional dimer interface (pLDDT >90) (residues 900-1032) (**Figure 3B, Supplementary Figure S5**) (21). Supporting this model, we observe that the isolated MORC2 C-terminus (residues 900-1032) forms a dimer in solution (**Figure 3C**). These data indicate that MORC2 contains N- and C-terminal dimerization interfaces like other GHL-type ATPases.

**Figure 3.**
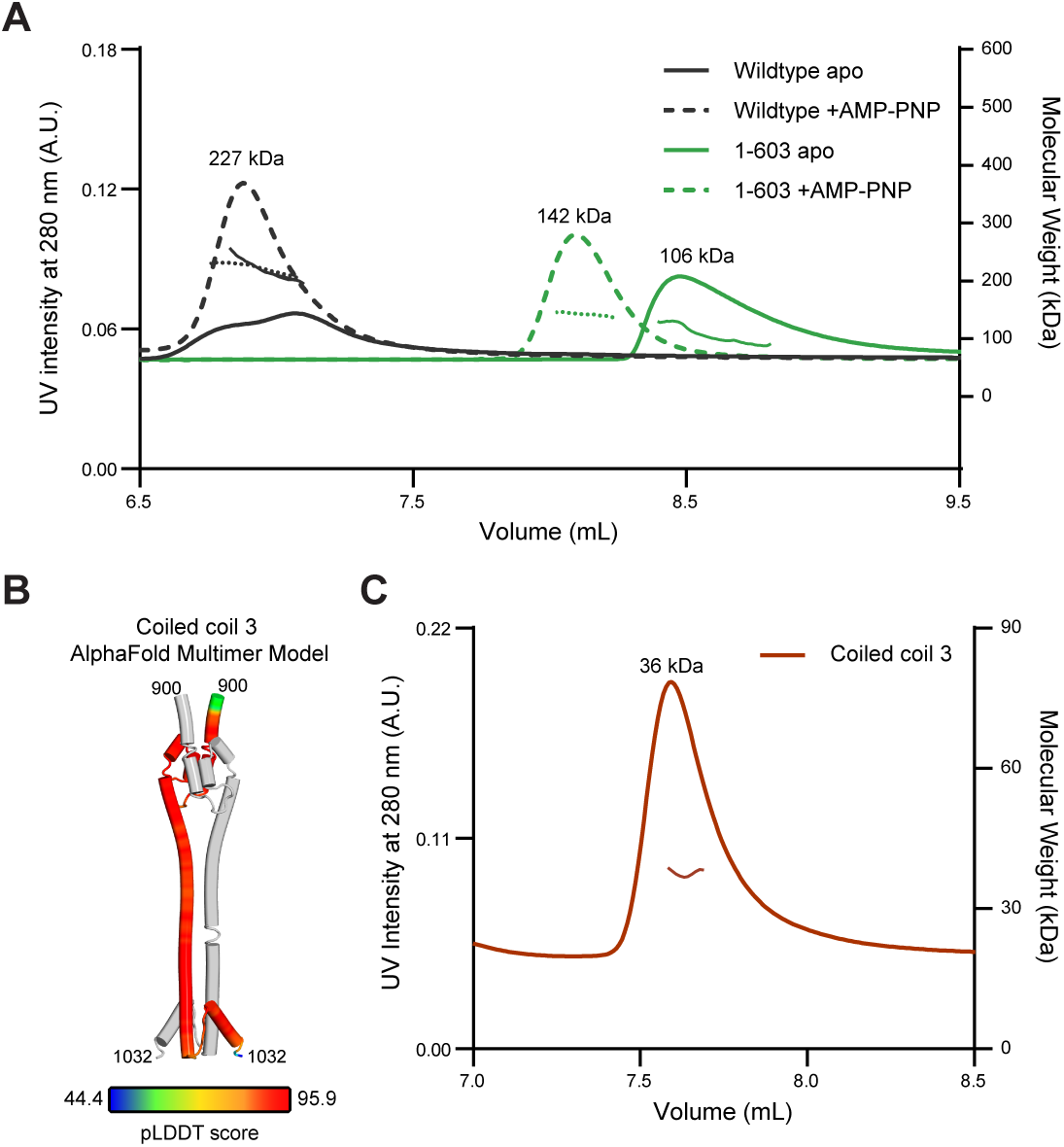
MORC2 homodimerizes via two distinct interfaces. **A.** MORC2 dimerizes in the presence and absence of AMP-PNP. Size exclusion chromatography coupled to multi-angle light scattering (SEC-MALS) experiments with wildtype (monomer MW =117 kDa) and 1-603 (monomer MW =70 kDa) MORC2 in the presence or absence of 1 mM AMP-PNP. 2 mg/mL of MORC2 was applied to a WC-030 column (Wyatt Technology) and elution was monitored by absorption at 280 nm (left y-axis). MALS analysis of molecular weight is shown on the right y-axis, and average molecular weight calculations are shown across the center of each peak. **B.** AlphaFold Multimer model of coiled coil 3 domain dimer. Chain A is colored grey and chain B is colored by pLDDT score. **C.** MORC2 coiled coil 3 dimerizes in solution. SEC-MALS experiment performed with 1.5 mg/mL of purified MORC2 coiled coil 3 (monomer MW =16 kDa) on a WC-010 column (Wyatt Technology). Elution was monitored by absorption at 280 nm (left y-axis). MALS analysis of molecular weight is shown on the right y-axis, and the average molecular calculation is shown across the center of the peak.

### MORC2 captures circular DNA substrates

Dimerization of GHL-type ATPases at their N- and C-termini can result in the formation of a central substrate binding lumen (61). To test whether MORC2 engages DNA via a central lumen, we utilized maltose binding protein (MBP)-tagged MORC2 in a pull-down assay with plasmid derived circular and linear DNA substrates (**Figure 4A**). If DNA is engaged inside a central binding lumen, it is expected that circular DNA will not be able to leave the protein when washed with high ionic strength solutions whereas linear DNA will be lost under these conditions because it is not topologically entrapped by the protein. MBP-MORC2 was incubated with linear or circular DNA and then added to amylose beads in the absence or presence of the ATP analog AMP-PNP. The amylose beads were washed extensively with either a low or high ionic strength containing solution (50 or 400 mM NaCl, respectively). MBP-MORC2 was eluted from the beads with maltose, digested with proteinase K, and the retained DNA was collected and separated by agarose gel electrophoresis. Circular DNA substrates are preferentially captured relative to linear substrates (∼30 fold) after a low ionic strength wash (**Figure 4B**). The addition of AMP-PNP does not substantially affect DNA retention under these conditions. In contrast, after a high ionic strength wash, retention of circular DNA substrates is enhanced with the inclusion of AMP-PNP, indicating that robust DNA retention requires the engagement of both dimerization interfaces. We also tested the ability of the aspartate DNA binding mutant and the ATP hydrolysis deficient E35A mutant to retain DNA (**Figure 4B**). The aspartate mutant captured almost no DNA under any condition. The E35A mutant retained circular substrates in the presence of AMP-PNP after low ionic strength washes. This effect was lost after high ionic strength washes, likely because the E35A GHKL domain dimerizes less stably (59). These results indicate that DNA binds within a central MORC2 lumen and that the DNA binding surface between residues 704-761 is required for this interaction.

**Figure 4.**
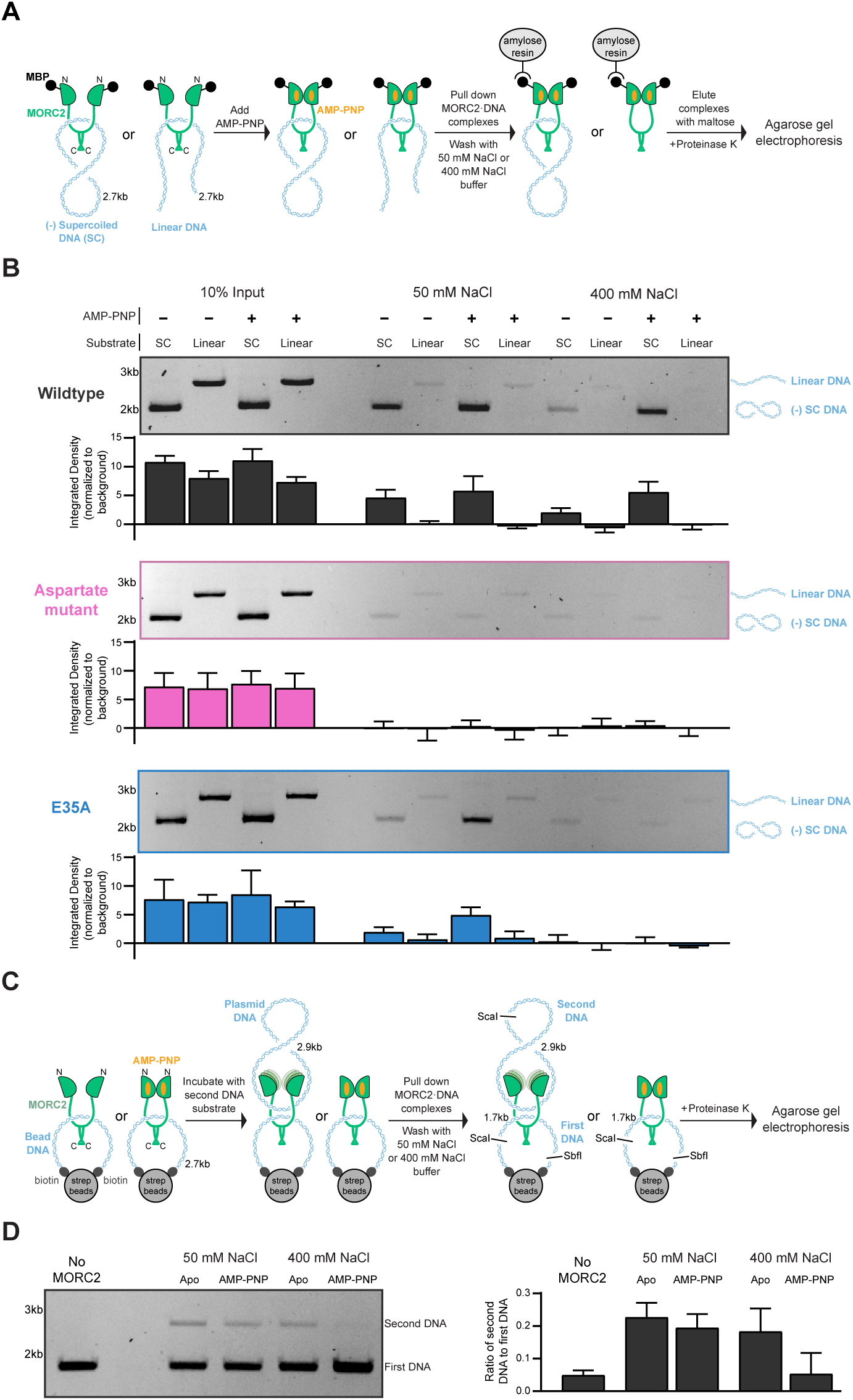
MORC2 can capture circular DNA substrates. **A.** Schematic of assay to assess MORC2 capture of linear or circular DNA substrates. **B.** N-terminally-maltose binding protein (MBP)-tagged wildtype, aspartate mutant, and E35A MORC2 (600 nM) were incubated with 100 nM supercoiled or linear pUC19. Samples were then added to amylose resin and washed in either low salt (50 mM NaCl) or high salt (400 mM NaCl) containing buffers before eluting the samples from the beads with maltose. Eluted samples were treated with proteinase K. DNA was resolved on a 1% (w/v) TAE agarose gel. Quantification of the band intensity from input and retained DNA bands normalized to the background are shown below each gel. Error bars correspond to the standard deviation between three replicate experiments. **C.** Schematic of assay to assess MORC2 association with two circular DNA substrates. **D.** A biotin tagged DNA was conjugated to streptavidin magnetic beads to create a pseudo circular substrate. MORC2 (600 nM) was incubated with 20 µL of the beads in the presence or absence of 1 mM AMP-PNP. Supercoiled pBlueScript plasmid DNA (200 nM) was added before washing the beads with either low salt (50 mM NaCl) or high salt (400 mM NaCl) containing buffer. Samples were resuspended in 1X CutSmart buffer (New England Biolabs). DNA was released from the beads by digestion with ScaI and SbfI at 37°C for 1 hour before proteinase K treatment. DNA was resolved on a 1% (w/v) TAE agarose gel. The intensity of the first DNA and second DNA substrate bands were quantified, normalized to the background, and are presented as a ratio of second DNA: first DNA band intensity. Error bars correspond to the standard deviation between three replicate experiments.

### MORC2 can bridge two DNA substrates

MORC family proteins are proposed to compact DNA by engaging multiple strands of DNA simultaneously (15). We assessed if MORC2 can bind multiple DNA strands using a dual DNA pulldown assay (**Figure 4C**). A DNA substrate containing ScaI and Sbf1 restriction sites was biotinylated on both ends and applied to streptavidin-coated magnetic beads to form a pseudo-circular DNA substrate. MORC2 was incubated with the pseudo-circular DNA substrate, and AMP-PNP was added or omitted to either stabilize the N-terminal dimerization interface or allow for its transient opening. Next, a second, circular plasmid DNA substrate with a single ScaI restriction site was added, and the samples were washed with either low or high ionic strength containing solutions. DNA was released from the beads by ScaI and Sbf1 restriction enzyme digest resulting in 1.7 kb and 2.9 kb DNA fragments corresponding to the biotinylated DNA and plasmid DNA species, respectively. MORC2 was digested with proteinase K, and the resulting DNA was collected and separated by agarose gel electrophoresis.

Control experiments without MORC2 show no retention of the second plasmid substrate. Under low ionic strength conditions, MORC2 associates with both DNA substrates efficiently in an AMP-PNP independent manner (**Figure 4D**). A high ionic strength wash, however, shows that MORC2 cannot effectively associate with the second plasmid DNA substrate when AMP-PNP is present (**Figure 4D**). These data indicate that MORC2 can capture two or more circular DNA substrates simultaneously and that stable, second strand capture requires the opening of the N-terminal GHKL dimerization interface.

### Phosphorylation inhibits MORC2 nuclear localization and mitotic chromosome association

We next explored how our biochemically identified phosphorylation and DNA binding sites in the MORC2 C-terminus affect MORC2 function in cells. We first investigated how MORC2 phosphorylation affects MORC2 cellular localization. To do this, we introduced EGFP tagged MORC2 ectopically into HeLa cells using the piggyBac transposase system (**Methods**). MORC2 expression was induced with doxycycline for 48 hours. Wildtype MORC2 and phosphodead MORC2, where phosphorylation sites are substituted with alanine, are localized to the nucleus (**Figure 5A**). In contrast, phosphomimetic MORC2, where phosphorylation sites are mutated to aspartate or glutamate, is found in the cytoplasm (**Figure 5A**). A predicted nuclear localization sequence (NLS) lies between the phosphorylation sites, corresponding to MORC2 residues 755-762 (**Figure 5B, Supplementary Figure S7A**) (62). Indeed, EGFP attached to MORC2 residues 734-771 localizes to the nucleus (**Figure 5A**). We note that our aspartate DNA binding mutant contains substitutions in the predicted NLS and localizes to the cytoplasm, further supporting the identified NLS (**Supplementary Figure S7B**). These results suggest that phosphorylation of MORC2 masks its NLS and prevents MORC2 from being localized to the nucleus.

**Figure 5.**
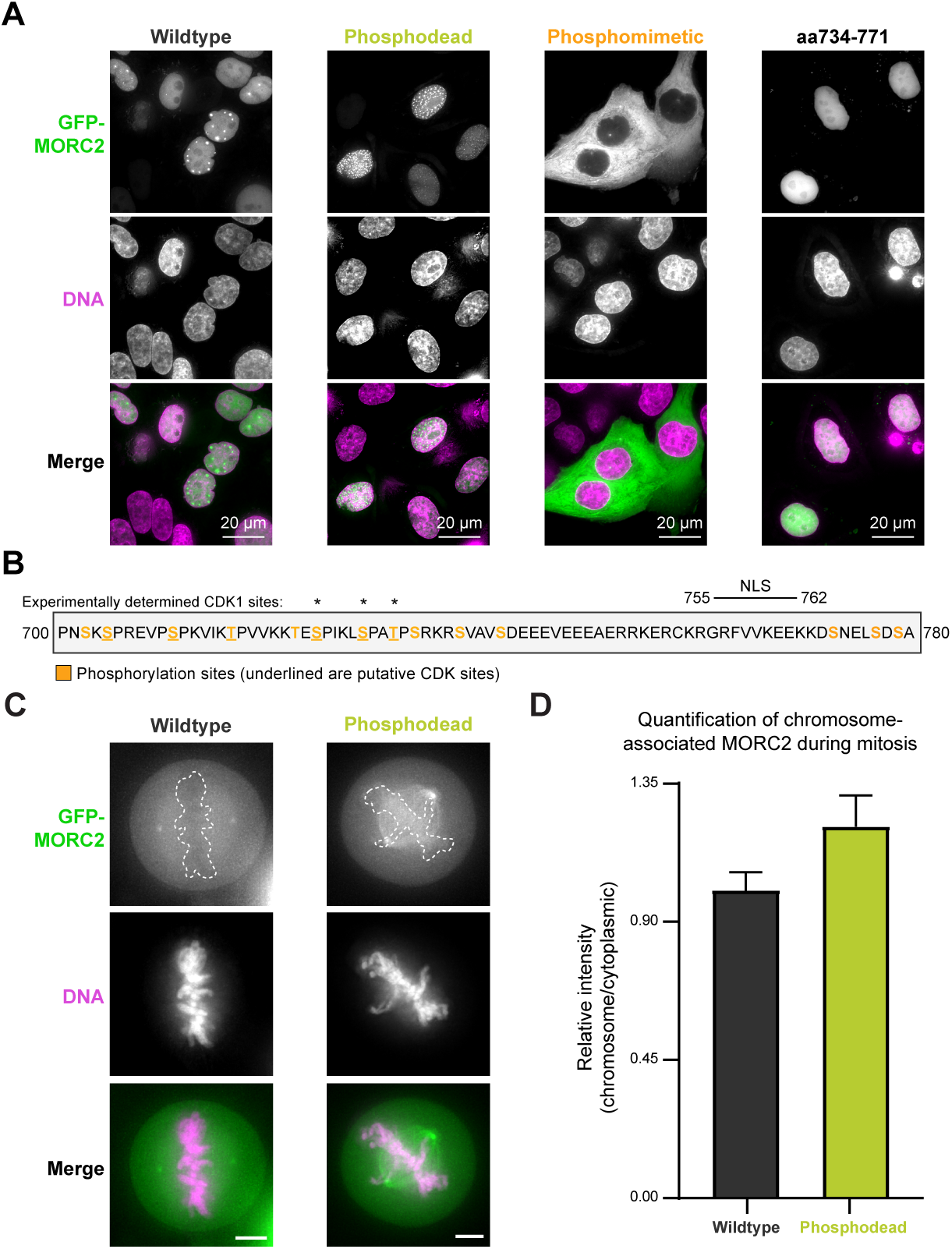
Phosphorylation regulates MORC2 localization and DNA association in cells. **A.** Effect of phosphorylation mutations on MORC2 localization in interphase HeLa cells. Representative confocal microscopy images of interphase HeLa cells overexpressing EGFP- wildtype MORC2, EGFP- phosphodead MORC2, EGFP- phosphomimetic MORC2, and EGFP- MORC2 734-771. **B.** Schematic of MORC2 domain encompassing the predicted NLS and phosphorylation sites mutated in the phosphomimetic mutant. Experimentally determined CDK1 phosphorylation sites are marked with asterisks. Putative CDK sites are underlined. **C.** Effect of phosphorylation on MORC2 chromosome localization during mitosis. Representative confocal microscopy images of mitotic HeLa cells overexpressing EGFP- wildtype MORC2 or EGFP- phosphodead MORC2. Scale bar represents 5 µm. Dotted lines are drawn around the chromosomes. **D.** Quantification of chromosome associated MORC2 during mitosis. The ratio of the mean fluorescence intensity of chromosome associated protein to the mean fluorescence intensity of the cytoplasm, relative to wildtype, is shown (**Methods**). Error bars correspond to the standard deviation between individual cells (wildtype n=19 and phosphodead n=20).

MORC2 is a known substrate of cyclin-dependent kinase 1 (CDK1) during mitosis (63), and nine of the seventeen MORC2 phosphorylation sites we detect contain the CDK1 motif Ser/Thr-Pro, where Ser/Thr is phosphorylated. We thus monitored the localization of wildtype and phosphodead MORC2 during mitosis. We observe that wildtype is largely excluded from chromosomes, whereas phosphodead MORC2 retains some association with mitotic chromosomes. Specifically, EGFP-phosphodead MORC2 chromosome:cytoplasmic intensity is 21% higher than EGFP-wildtype MORC2 chromosome:cytoplasmic intensity (**Figure 5C and Figure 5D**), suggesting that preventing MORC2 phosphorylation limits its eviction from mitotic chromosomes. Together, phosphorylation serves at least two purposes for MORC2 regulation in cells: 1) it blocks MORC2 localization to the nucleus and 2) supports MORC2 eviction from chromosomes during mitosis, likely by reducing the affinity of MORC2 for DNA.

### The C-terminal DNA binding region is required for MORC2 gene silencing in cells

We next tested how the MORC2 DNA binding region affects MORC2’s silencing function in cells. We appended an artificial SV40 NLS sequence (35) to EGFP, EGFP-wildtype MORC2, and the EGFP-MORC2 DNA binding deficient aspartate mutant to ensure all MORC2 proteins were localized to the nucleus (**Supplementary Figure S7C),** and ectopically introduced these constructs into a MORC2 knockout HeLa cell line (**Methods, Supplementary Figure S8**). Doxycycline was added to induce EGFP or EGFP-MORC2 expression for 48 hours, and total RNA was collected for spike-in normalized RNA sequencing (**Methods**). We confirmed that expression levels of MORC2 were similar between the wildtype and aspartate mutant samples (**Supplementary Figure S9B**). Analysis of human protein-coding genes shows that expression of wildtype EGFP-MORC2 leads to significant upregulation of 69 genes and significant downregulation of 197 genes in comparison to control cells expressing EGFP (**Figure 6A**). Almost 40% of the genes downregulated by wildtype MORC2 are intronless or contain exons longer than 1 kb, of which 26 are zinc finger (ZNF) genes, consistent with prior studies (**Supplementary Figure S9E**) (5). In contrast, expression of EGFP-tagged aspartate mutant MORC2 leads to significant upregulation of 4 genes and significant downregulation of 13 genes in comparison to control cells expressing EGFP (**Figure 6A**). We compared the fold repression of intronless or long-exon containing genes that are significantly repressed by wildtype MORC2 with their fold repression after expression of the aspartate mutant, and the aspartate mutant exhibits weaker repression for almost all genes analyzed (**Figure 6B**).

**Figure 6.**
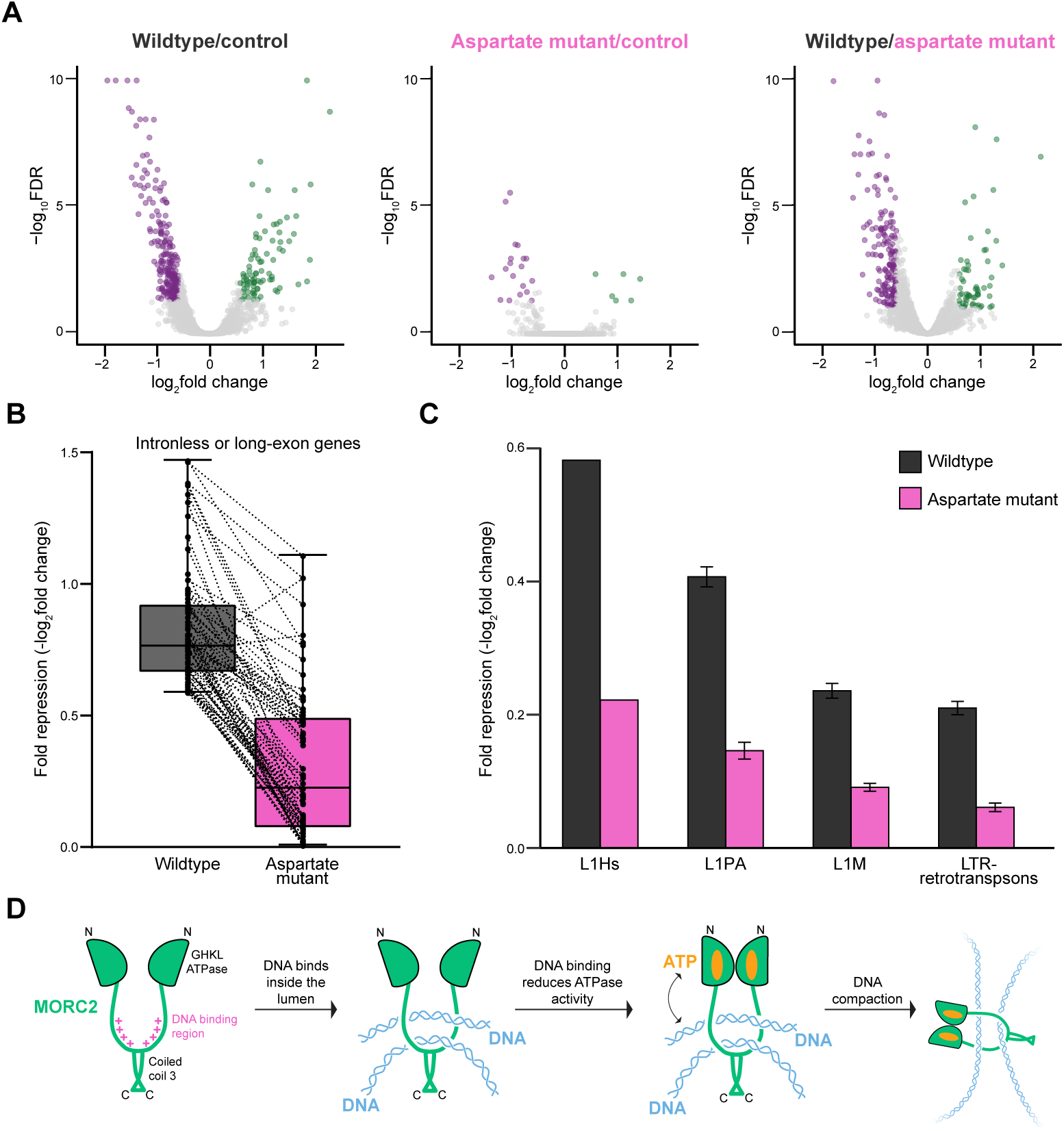
DNA binding regulates MORC2 gene silencing activity in cells. **A.** Volcano plots of RNAseq reads for overexpression of wildtype MORC2 versus control, overexpression of aspartate mutant MORC2 versus control, and wildtype MORC2 versus aspartate mutant MORC2 with spike normalization (**Methods**). Significant upregulated genes are shown in green and significant downregulated genes are shown in purple from three biological replicates. Significant genes are classified as those that meet the fold change > 1.5 and FDR > 0.05 cutoffs. **B.** Fold repression (-log_2_fold change) analysis of intronless and long-exon containing genes exhibiting a significant degree of repression after overexpression of wildtype MORC2 (n=75) in comparison to their fold repression after overexpression of aspartate mutant MORC2. Error bars correspond to the minimum and maximum values and the central bar represents the median value. Dotted lines connect genes between the two conditions. **C.** Average fold repression (-log_2_fold change) of retrotransposon subfamilies as measured by RNAseq in cells with overexpression of wildtype or aspartate mutant MORC2 versus control cells using the TEtranscripts tool. Error bars represent the standard error between the elements in each subfamily (L1Hs n = 1, L1PA n = 30, L1M n = 86, and LTR-retrotransposons n = 347) **D.** Model of how MORC2 engages DNA to promote compaction. MORC2 contains a DNA binding region between the GHKL domain and coiled coil 3 domain dimerization interfaces. DNA binding inside the lumen of the dimer communicates to the ATPase domain, to favor an ATP-bound homodimer conformation that is incompatible with ATP hydrolysis. MORC2 association with multiple DNA segments may allow MORC2 to bridge distal regions of DNA to contribute to compaction.

We next assessed the role of MORC2 in silencing retrotransposons using the TEtranscripts tool, which improves mapping of transposable elements (48). Expression of wildtype MORC2 represses evolutionarily young retrotransposons as previously observed (28, 32) whereas expression of the DNA binding deficient aspartate mutant weakly represses expression of retrotransposons (**Figure 6C**). The most highly repressed retrotransposons are the youngest and second youngest LINE1 classes in the human genome, L1Hs and L1PA, respectively. Wildtype MORC2 weakly represses expression of older LINE1s from the L1M class and LTR-retrotransposons. Taken together, these results indicate that the MORC2 C-terminal DNA binding region is required for silencing known MORC2 targets.

## Discussion

Here we have identified a MORC2 DNA binding region (residues 704-761) that lies in between its GHKL domain and C-terminal domain dimerization interfaces. This region is required for the capture of circular DNA substrates by MORC2, indicating that MORC2 can topologically entrap DNA. Notably, DNA binding reduces MORC2 ATP hydrolysis activity. We also observe that MORC2 is heavily phosphorylated, and this phosphorylation appears to regulate MORC2 cellular localization and association with DNA. Finally, we show that the identified DNA binding region is required for MORC2 gene silencing in HeLa cells.

Our data integrated with previous results provide a model for how MORC2 could promote chromatin compaction and gene silencing (**Figure 6D**). In this model, transient opening of the MORC2 GHKL domain would allow for DNA substrates to enter the MORC2 lumen and associate with the MORC2 C-terminal DNA binding region. DNA binding is communicated to the GHKL domain to presumably favor an ATP-bound GHKL homodimer conformation that is incompatible with ATP hydrolysis. This would serve to topologically clamp MORC2 around DNA, by prolonging the ATP-bound homodimerized state of the GHKL domain. The mechanism for coupling between the ATPase and C-terminal DNA binding region is presently unclear. The concept that ATPase rate and MORC2 silencing activity are coupled is borne out in neuropathic disease associated MORC2 mutants that alter GHKL dimerization efficiency and ATP hydrolysis rates (18). Specifically, MORC2 S87L stabilizes GHKL dimerization and has negligible ATPase activity and high levels of silencing in cells (18). Conversely, MORC2 T424R reduces GHKL dimerization, has high ATPase levels, and reduced levels of silencing compared to the wildtype protein (18). Thus, ATP hydrolysis rate appears to be inversely associated with MORC2 silencing activity in cells, and our results indicate that DNA binding leads to reduced ATPase activity. Finally, association of MORC2 with multiple DNA segments, as we observe here, could lead to compaction of the DNA by bringing distal DNA segments together and thereby promote gene silencing.

These results support a general mechanism for how MORC family proteins engage with DNA to promote gene silencing. All MORC family proteins contain an N-terminal GHKL domain and a predicted C-terminal dimerization domain (1, 16). Dimerization of MORC proteins around DNA at both interfaces could result in topological entrapment of chromatin substrates. Indeed, *C. elegans* MORC-1 and human MORC2 appear to topologically engage with DNA (15). It is not yet known if MORC3 and MORC4 can topologically entrap DNA, and if the DNA binding surface we identified on MORC2 extends to the other MORC family members. From our work, it is also unclear how the previously identified MORC2 coiled coil 1 DNA interaction is used to promote gene silencing (18). This domain is absent in MORC3 and MORC4. It is possible that the coiled coil 1 interface is required for localization but not for topological entrapment of DNA substrates.

MORC family proteins are localized to unique genomic positions. Our results suggest that MORC2 does not bind DNA in a sequence specific manner. Rather than recognizing a specific motif, MORC2 can be directed to retroelements and intronless genes, in part, through its association with the Human Silencing Hub (HUSH) complex (5). In addition to protein complexes, histone post-translational modifications can be used to localize MORC proteins to specific genomic regions. Histone H3 trimethylation of lysine 4 is associated with MORC3 and MORC4 localization, driven through an interaction with the modified histone tail and the MORC CW domain (8). MORC1 and MORC2 do not engage with histone proteins in a specific manner, indicating their recruitment to specific genomic regions is likely independent of chromatin modification state (33). Finally, post-translational modifications of MORC proteins can drive their localization to specific cellular locations, as observed here with phosphorylation of MORC2 to limit MORC2 interactions with chromatin or as previously observed with MORC3 SUMOlyation that serves to localize MORC3 to PML bodies (3). Future work with full-length MORC proteins will uncover whether all members use a similar mechanism as MORC2 to associate with chromatin and define how MORC family proteins are localized to specific genomic locations to induce silencing.

## Acknowledgements

We would like to thank Iain Cheeseman for supporting the cell-based experiments, Greg Dodge for SEC-MALS assistance, and the past and present members of the Vos lab for their support. We thank Eric Spooner at the Whitehead Institute for Biomedical Research Proteomics Core facility and Harvard Medical School Taplin Mass Spectrometry Facility for protein mass spectrometry analysis, the MIT BioMicro Center and Stuart Levine for their assistance with RNA sequencing, and the MIT Biophysical Instrumentation Facility for use of the CD spectrometer. We would further like to thank Monika Raabe from the Bioanalytical Mass Spectrometry Group at the Max Planck Institute for Multidisciplinary Sciences in Göttingen for the experimental and technical support in crosslinking mass spectrometry experiments.

## Data availability

Raw data for RNA sequencing is available under Gene Expression Omnibus [accession code GSE262311]. The mass spectrometry proteomics data have been deposited to the ProteomeXchange Consortium via the PRIDE partner repository [dataset identifier PXD050648]. All reagents generated in this study are available upon reasonable request and will be fulfilled by the corresponding author (SMV).

