## Supplementary material for "Identification and characterization of a human MORC2 DNA binding region that is required for gene silencing": SI Material

**SI Table 1. Identity of contaminant band (Fig1B) as assessed by mass spectrometry.**

| Species: *Trichoplusia ni* | | | |
|---|---|---|---|
| **Unique Counts** | **Total Counts** | **Protein Name** | **Gene ID** |
| 89 | 95 | Serine/threonine-protein kinase TOR | LOC113507625 |
| 83 | 91 | acetyl-CoA carboxylase | LOC113498903 |
| 71 | 76 | DmX-like protein 2 isoform X1 | LOC113503468 |
| 68 | 74 | Talin-1 isoform X4 | LOC113499982 |
| 65 | 70 | Triple functional domain protein isoform X1 | LOC113504138 |
| 62 | 71 | Dedicator of cytokinesis protein 7 isoform X3 | LOC113501546 |
| 61 | 65 | ubiquitinyl hydrolase 1 | LOC113492721 |
| 59 | 63 | U5 small nuclear ribonucleoprotein 200 kDa helicase | LOC113505650 |
| 55 | 62 | Beta-galactosidase | LOC113504558 |
| 53 | 55 | LOW QUALITY PROTEIN: nuclear pore complex protein Nup205 | LOC113493667 |
| 47 | 52 | Transcription elongation factor spt6 | LOC113502227 |
| 44 | 48 | CCR4-NOT transcription complex subunit 1 isoform X3 | LOC113504324 |
| 41 | 43 | Fatty acid synthase isoform X1 | LOC113498344 |
| 39 | 43 | Pre-mRNA-processing-splicing factor 8 | LOC113502029 |
| 35 | 39 | WD repeat and FYVE domain-containing protein 3 isoform X3 | LOC113497332 |
| 34 | 36 | Protein 4.1 homolog isoform X1 | LOC113494423 |
| 34 | 36 | Maternal protein tudor-like isoform X1 | LOC113498000 |
| 34 | 36 | Uncharacterized protein LOC113503664 | LOC113503664 |
| 30 | 31 | LOW QUALITY PROTEIN: baculoviral IAP repeat-containing protein 6-like | LOC113492500 |

| Species: *Homo sapians* | | | |
|---|---|---|---|
| **Unique Counts** | **Total Counts** | **Protein Name** | **Gene ID** |
| 95 | 305 | ATPase MORC2 | MORC2 |

**SI Table 2. $K_{d,app}$ values for all proteins, substrates, and conditions tested.**

| **Protein** | **Substrate** | **Condition** | **$K_{d,app}$ (nM)** |
|---|---|---|---|
| Dephosphorylated MORC2 | 35mer random dsDNA | apo | 17 ± 4 |
| Phosphorylated MORC2 | 35mer random dsDNA | apo | 285 ± 83 |
| Phosphodead | 35mer random dsDNA | apo | 25 ± 3 |
| Phosphomimetic | 35mer random dsDNA | apo | 512 ± 147 |
| 1-603 | 35mer random dsDNA | apo | 446 ± 119 |
| Aspartate mutant | 35mer random dsDNA | apo | 380± 120 |
| Alanine mutant | 35mer random dsDNA | apo | 350± 60 |
| Subset N | 35mer random dsDNA | apo | 90 ± 20 |
| Subset C | 35mer random dsDNA | apo | 60 ± 20 |
| Wildtype | 35mer random dsDNA | apo | 5 or less |
| Wildtype | 149 bp Widom 601 dsDNA | apo | 5 or less |
| Wildtype | Nucleosome | apo | 16 ± 2 |
| Wildtype | 35mer ssDNA | apo | 51 ± 9 |
| Wildtype | 35mer ssRNA | apo | 49 ± 6 |
| Wildtype | 35mer high GC | apo | 5 or less |
| Wildtype | 35mer high AT | apo | 5 or less |
| Wildtype | 25mer random dsDNA | apo | 21 ± 2 |
| Wildtype | 45mer random dsDNA | apo | 11 ± 3 |
| Wildtype | 65mer random dsDNA | apo | 5 or less |
| Wildtype | 35mer random dsDNA | AMP-PNP | 30 ± 5 |
| Wildtype | 35mer random dsDNA | ADP | 36 ± 8 |
| Wildtype | 35mer random dsDNA | ATP | 86 ± 15 |
| Wildtype | 35mer random dsDNA | ATPγS | 275 ± 51 |

### SI Table 3. Mechlorethamine and UV-induced protein-DNA crosslinking sites.

*UV-induced crosslinking sites*

| index | RT | precursor m/z | score | charge | sequence | start | end | NuXL:NA | NuXL:NT | NuXL:best_localization | NuXL:best_localization_position | NuXL:best_localization_score | precursor_mz_error_ppm |
|---|---|---|---|---|---|---|---|---|---|---|---|---|---|
| 10033 | 1512 | 836.351318 | 7.98E-06 | 2 | AM(Oxidation)GEHLAQYWK | 442 | 452 | T | T | AMgEHLAQYWK | 2 | 0.46096 | 4.82232659 |
| 21987 | 3187 | 1033.88635 | 0.003992589 | 2 | DIQMAETSPEGTK | 222 | 234 | AG | A | DIQMAETSPEGTk | 12 | 0.08188 | -0.9964457 |
| 8647 | 1326 | 608.2481 | 0.003992664 | 4 | DLDGM(Oxidation)FIYNCSRLIK | 378 | 392 | TT | T | DLDGMFIyNCSRLIK | 7 | 0.67426 | -3.6523695 |
| 12616 | 1858 | 823.322571 | 0.007701015 | 2 | EDTM(Oxidation)TCLFLSR | 122 | 132 | A-H3N1 | A | EDTMTCLfLSR | 7 | 0.72102 | -0.4642795 |
| 9392 | 1425 | 961.8713 | 7.62E-06 | 2 | FDYVPTDTTPR | 828 | 838 | CT | C | FDYVpTDTTpR | 9 | 0.05199 | 5.60342229 |
| 9267 | 1408 | 961.8706 | 0.003992478 | 2 | FDYVPTDTTPR | 828 | 838 | CT | C | FDYVpTDTTpR | 9 | 0.06368 | 4.87567012 |
| 8978 | 1369 | 817.347229 | 7.07E-06 | 2 | FDYVPTDTTPR | 828 | 838 | T | T | FDyVPTDTTPR | 2 | 1.01463 | 5.51259717 |
| 9834 | 1485 | 817.346619 | 7.06E-06 | 2 | FDYVPTDTTPR | 828 | 838 | T | T | FDyVPTDTTPR | 2 | 1.02506 | 4.76584609 |
| 9779 | 1477 | 817.347412 | 7.06E-06 | 2 | FDYVPTDTTPR | 828 | 838 | T | T | FDyVPTDTTPR | 2 | 1.04658 | 5.7366225 |
| 3540 | 659.8 | 757.31311 | 7.34E-06 | 2 | FVVKEEK | 764 | 770 | AT | A | FVVKEEK | -1 | 0 | 4.31315599 |
| 4349 | 771.5 | 505.210999 | 8.26E-06 | 3 | FVVKEEK | 764 | 770 | AT | A | FVVKEEK | -1 | 0 | 3.97936082 |
| 4226 | 754.7 | 765.311584 | 7.96E-06 | 2 | FVVKEEK | 764 | 770 | GT | G | FVVKEEK | -1 | 0 | 5.59646638 |
| 4713 | 820.5 | 765.310852 | 8.25E-06 | 2 | FVVKEEK | 764 | 770 | GT | G | FVVKEEK | -1 | 0 | 4.63943661 |
| 4085 | 735.1 | 600.781738 | 7.34E-06 | 2 | FVVKEEK | 764 | 770 | T | T | FVVKEEK | -1 | 0 | 1.1614241 |
| 4806 | 833.2 | 752.807495 | 6.14E-06 | 2 | FVVKEEK | 764 | 770 | TT | T | FVVKEEK | -1 | 0 | 4.56246897 |
| 4471 | 788.2 | 757.314087 | 7.10E-06 | 2 | FVVKEEK | 764 | 770 | AT | A | FVVKeeK | 5 | 0.00707 | 5.60267112 |
| 4570 | 801.5 | 757.313904 | 7.09E-06 | 2 | FVVKEEK | 764 | 770 | AT | A | FVVKeeK | 5 | 0.00941 | 5.36088703 |
| 4075 | 733.7 | 600.7842 | 7.40E-06 | 2 | FVVKEEK | 764 | 770 | T | T | fVVKEEK | 0 | 0.02402 | 5.25895479 |
| 4457 | 786.3 | 765.311218 | 7.40E-06 | 2 | FVVKEEK | 764 | 770 | GT | G | FVVkEEk | 6 | 0.02593 | 5.1179515 |

| | | | | | | | | | | | | | |
|---|---|---|---|---|---|---|---|---|---|---|---|---|---|
| 5093 | 871.7 | 752.808044 | 7.34E-06 | 2 | FVVKEEK | 764 | 770 | TT | T | FvVKEEK | 1 | 0.03597 | 5.2921627 |
| 4134 | 741.8 | 765.311157 | 0.003992613 | 2 | FVVKEEK | 764 | 770 | GT | G | FVVkEEk | 6 | 0.03685 | 5.03819901 |
| 3684 | 680 | 745.307678 | 7.34E-06 | 2 | FVVKEEK | 764 | 770 | CT | C | FvVKEEK | 1 | 0.07419 | 4.63001075 |
| 4054 | 730.8 | 745.307129 | 7.99E-06 | 2 | FVVKEEK | 764 | 770 | CT | C | FvVKEEK | 1 | 0.11419 | 3.89297427 |
| 3975 | 720.1 | 745.3083 | 6.70E-06 | 2 | FVVKEEK | 764 | 770 | CT | C | FvVKEEK | 1 | 0.18691 | 5.46427053 |
| 3903 | 710.2 | 745.307129 | 6.80E-06 | 2 | FVVKEEK | 764 | 770 | CT | C | FvVKEEK | 1 | 0.18979 | 3.89297427 |
| 3444 | 646.5 | 745.307678 | 7.34E-06 | 2 | FVVKEEK | 764 | 770 | CT | C | FvVKEEK | 1 | 0.33493 | 4.63001075 |
| 3378 | 637.2 | 745.307434 | 6.16E-06 | 2 | FVVKEEK | 764 | 770 | CT | C | FvVKEEK | 1 | 0.37333 | 4.30243898 |
| 4004 | 724.1 | 600.783936 | 4.45E-06 | 2 | FVVKEEK | 764 | 770 | T | T | fVVKEEK | 0 | 0.37715 | 4.81877258 |
| 3873 | 706 | 670.7843 | 0.003992545 | 2 | GRFVVK | 762 | 767 | AT | A | GRFVVK | -1 | 0 | 4.83648033 |
| 4026 | 727.1 | 670.784058 | 7.30E-06 | 2 | GRFVVK | 762 | 767 | AT | A | GRFVVK | -1 | 0 | 4.47513617 |
| 4184 | 748.8 | 670.784 | 7.91E-06 | 2 | GRFVVK | 762 | 767 | AT | A | GRFVVK | -1 | 0 | 4.3892405 |
| 4250 | 757.9 | 670.7846 | 8.18E-06 | 2 | GRFVVK | 762 | 767 | AT | A | GRFVVK | -1 | 0 | 5.28372015 |
| 4431 | 782.7 | 670.784546 | 7.30E-06 | 2 | GRFVVK | 762 | 767 | AT | A | GRFVVK | -1 | 0 | 5.20306557 |
| 4519 | 794.7 | 670.783997 | 7.30E-06 | 2 | GRFVVK | 762 | 767 | AT | A | GRFVVK | -1 | 0 | 4.384145 |
| 4582 | 803.1 | 670.783997 | 6.99E-06 | 2 | GRFVVK | 762 | 767 | AT | A | GRFVVK | -1 | 0 | 4.384145 |
| 3728 | 685.9 | 658.777893 | 6.65E-06 | 2 | GRFVVK | 762 | 767 | CT | C | GRFVVK | -1 | 0 | 3.72476817 |
| 4294 | 763.9 | 658.7786 | 8.18E-06 | 2 | GRFVVK | 762 | 767 | CT | C | GRFVVK | -1 | 0 | 4.7978707 |
| 3942 | 715.5 | 678.781189 | 0.003992549 | 2 | GRFVVK | 762 | 767 | GT | G | GRFVVK | -1 | 0 | 3.94191061 |
| 4077 | 734 | 678.780762 | 7.30E-06 | 2 | GRFVVK | 762 | 767 | GT | G | GRFVVK | -1 | 0 | 3.31247688 |
| 4147 | 743.7 | 678.7816 | 7.61E-06 | 2 | GRFVVK | 762 | 767 | GT | G | GRFVVK | -1 | 0 | 4.54746183 |
| 4210 | 752.5 | 678.781433 | 7.61E-06 | 2 | GRFVVK | 762 | 767 | GT | G | GRFVVK | -1 | 0 | 4.30158703 |

| | | | | | | | | | | | | | |
|---|---|---|---|---|---|---|---|---|---|---|---|---|---|
| 4272 | 761 | 678.781433 | 0.003992545 | 2 | GRFVVK | 762 | 767 | GT | G | GRFVVK | -1 | 0 | 4.30158703 |
| 4611 | 807 | 678.781189 | 8.20E-06 | 2 | GRFVVK | 762 | 767 | GT | G | GRFVVK | -1 | 0 | 3.94191061 |
| 4674 | 815.3 | 678.782532 | 6.98E-06 | 2 | GRFVVK | 762 | 767 | GT | G | GRFVVK | -1 | 0 | 5.92013091 |
| 4731 | 823 | 678.781494 | 7.30E-06 | 2 | GRFVVK | 762 | 767 | GT | G | GRFVVK | -1 | 0 | 4.39150613 |
| 5126 | 875.9 | 678.781128 | 7.61E-06 | 2 | GRFVVK | 762 | 767 | GT | G | GRFVVK | -1 | 0 | 3.8519915 |
| 5385 | 910.8 | 666.279053 | 7.91E-06 | 2 | GRFVVK | 762 | 767 | TT | T | GRFvVK | 3 | 0.00352 | 5.67398786 |
| 4127 | 740.8 | 658.778503 | 6.65E-06 | 2 | GRFVVK | 762 | 767 | CT | C | gRFVVK | 0 | 0.00612 | 4.65126227 |
| 4669 | 814.7 | 670.783691 | 7.88E-06 | 2 | GRFVVK | 762 | 767 | AT | A | GRfVVK | 2 | 0.01262 | 3.92918912 |
| 4614 | 807.4 | 666.27771 | 6.99E-06 | 2 | GRFVVK | 762 | 767 | TT | T | GRFVvK | 4 | 0.01607 | 3.65864451 |
| 3809 | 697.3 | 658.777832 | 8.18E-06 | 2 | GRFVVK | 762 | 767 | CT | C | GRfVVK | 2 | 0.01785 | 3.63211876 |
| 4064 | 732.2 | 658.77832 | 7.61E-06 | 2 | GRFVVK | 762 | 767 | CT | C | gRFVVK | 0 | 0.02852 | 4.37331404 |
| 4461 | 786.8 | 666.278625 | 7.30E-06 | 2 | GRFVVK | 762 | 767 | TT | T | GRFVvK | 4 | 0.05964 | 5.03274225 |
| 5299 | 899.2 | 666.27832 | 6.90E-06 | 2 | GRFVVK | 762 | 767 | TT | T | GrFVVK | 1 | 0.06294 | 4.57470967 |
| 4770 | 828.3 | 666.277893 | 6.33E-06 | 2 | GRFVVK | 762 | 767 | TT | T | GRfVVK | 2 | 0.15849 | 3.93346406 |
| 3544 | 660.4 | 576.251465 | 8.34E-06 | 3 | GRFVVKEEK | 762 | 770 | AT | A | GRFVVKEEK | -1 | 0 | 2.80819964 |
| 3878 | 706.7 | 863.8746 | 7.81E-06 | 2 | GRFVVKEEK | 762 | 770 | AT | A | GRFVVKEEK | -1 | 0 | 4.01484035 |
| 3278 | 623.3 | 851.869568 | 8.06E-06 | 2 | GRFVVKEEK | 762 | 770 | CT | C | GRFVVKEEK | -1 | 0 | 4.75741264 |
| 3665 | 677.2 | 851.868652 | 8.35E-06 | 2 | GRFVVKEEK | 762 | 770 | CT | C | GRFVVKEEK | -1 | 0 | 3.68268039 |
| 3457 | 648.2 | 707.3444 | 8.34E-06 | 2 | GRFVVKEEK | 762 | 770 | T | T | GRFVVKEEK | -1 | 0 | 2.92890867 |
| 3814 | 697.9 | 576.2529 | 0.003992423 | 3 | GRFVVKEEK | 762 | 770 | AT | A | GRFvvKEEK | 4 | 0.00916 | 5.29871006 |
| 3456 | 648.1 | 851.869385 | 6.94E-06 | 2 | GRFVVKEEK | 762 | 770 | CT | C | GRFVVkEEK | 5 | 0.02006 | 4.54246619 |
| 3529 | 658.3 | 851.869751 | 6.23E-06 | 2 | GRFVVKEEK | 762 | 770 | CT | C | GRfVVKEEK | 2 | 0.03007 | 4.97235909 |

| | | | | | | | | | | | | | |
|---|---|---|---|---|---|---|---|---|---|---|---|---|---|
| 3904 | 710.4 | 871.872375 | 7.23E-06 | 2 | GRFVVKEEK | 762 | 770 | GT | G | GRfVVKEEK | 2 | 0.0409 | 4.34273353 |
| 4218 | 753.5 | 863.875366 | 0.003992421 | 2 | GRFVVKEEK | 762 | 770 | AT | A | GRFvvKEEK | 4 | 0.04651 | 4.90179086 |
| 3591 | 667 | 851.869507 | 8.09E-06 | 2 | GRFVVKEEK | 762 | 770 | CT | C | GRFvvKEEK | 4 | 0.05389 | 4.68576383 |
| 3631 | 672.5 | 863.8757 | 8.34E-06 | 2 | GRFVVKEEK | 762 | 770 | AT | A | GRfVVKEEK | 2 | 0.05545 | 5.28817842 |
| 4251 | 758 | 871.872253 | 7.23E-06 | 2 | GRFVVKEEK | 762 | 770 | GT | G | GRFvvKEEK | 4 | 0.06927 | 4.20272354 |
| 3844 | 702 | 851.8694 | 0.003992421 | 2 | GRFVVKEEK | 762 | 770 | CT | C | GRFvvKEEK | 4 | 0.07203 | 4.56034974 |
| 3828 | 699.9 | 871.87262 | 6.94E-06 | 2 | GRFVVKEEK | 762 | 770 | GT | G | GRfVVKEEK | 2 | 0.0751 | 4.62275351 |
| 3731 | 686.4 | 871.8737 | 7.47E-06 | 2 | GRFVVKEEK | 762 | 770 | GT | G | GRfVVKEEK | 2 | 0.08196 | 5.86189793 |
| 6219 | 1014 | 503.871521 | 0.003992573 | 3 | IFIHGHK | 255 | 261 | GG-H2O1 | G | IFiHGHK | 2 | 0.03428 | 4.18750337 |
| 13548 | 1986 | 1051.8924 | 0.003992565 | 2 | KEDTM(Oxidation)TCLFLSR | 121 | 132 | AG-H3N1 | A | KEDtMtCLFLSR | 5 | 0.11928 | -4.0837475 |
| 4398 | 778.2 | 546.754211 | 0.003992627 | 2 | KTESPIK | 722 | 728 | C-H3N1 | C | KtESPIK | 1 | 0.33112 | 3.52725315 |
| 14329 | 2093 | 844.824524 | 0.003992675 | 2 | PSTEEPVRR | 601 | 609 | TA-H3N1 | A | PsTEEPVRR | 1 | 0.12161 | -2.8379896 |
| 14402 | 2103 | 844.8244 | 0.00399267 | 2 | PSTEEPVRR | 601 | 609 | TA-H3N1 | A | PStEEPVRR | 2 | 0.17279 | -2.9846774 |
| 14991 | 2185 | 733.312317 | 0.00399262 | 2 | QLTEKIR | 563 | 569 | CC-H2O1 | C | QLTEkIR | 4 | 0.06764 | -4.2867793 |
| 7394 | 1163 | 662.9558 | 0.003992509 | 3 | QQQEKLEALQK | 570 | 580 | AA | A | qqqEKLEALqK | 9 | 0.176 | -2.7374137 |
| 7455 | 1171 | 744.868958 | 0.007700944 | 2 | QVQNRAITLR | 334 | 343 | C-H3N1 | C | qVQNRAITLR | 0 | 1.02484 | -0.8301039 |
| 11829 | 1752 | 992.063904 | 0.007700994 | 3 | QYEVGLQNLCNSYQSRADSR | 953 | 972 | AG-H3N1 | A | QYEVgLQNLCNSYQSRADSR | 4 | 0.15141 | 1.6019709 |
| 11479 | 1704 | 986.3979 | 7.58E-06 | 3 | QYEVGLQNLCNSYQSRADSR | 953 | 972 | TT | T | QYEVGLQNLCNsYQsRADsR | 18 | 0.1776 | -0.5892887 |
| 11583 | 1718 | 986.731567 | 0.007700994 | 3 | QYEVGLQNLCNSYQSRADSR | 953 | 972 | AA-H3N1 | A | QYEVGLQNLcNSYQSRADSR | 9 | 0.18494 | 0.90314665 |
| 11741 | 1740 | 992.062988 | 0.007700993 | 3 | QYEVGLQNLCNSYQSRADSR | 953 | 972 | AG-H3N1 | A | QYEVGlQNlCNSYQSRADSR | 8 | 0.20737 | 0.67911824 |
| 11675 | 1731 | 986.398 | 0.007700993 | 3 | QYEVGLQNLCNSYQSRADSR | 953 | 972 | TT | T | QYEVGLQNLCNsYQsRADsR | 18 | 0.2855 | -0.4879098 |
| 11401 | 1694 | 986.732727 | 7.34E-06 | 3 | QYEVGLQNLCNSYQSRADSR | 953 | 972 | AA-H3N1 | A | QYEVGLQnLCNSYQSRADSR | 7 | 0.45275 | 2.07840957 |

| | | | | | | | | | | | | | |
|---|---|---|---|---|---|---|---|---|---|---|---|---|---|
| 11045 | 1647 | 992.062683 | 0.007700988 | 3 | QYEVGLQNLCNSYQSRADSR | 953 | 972 | AG-H3N1 | A | QYEVGlQNlCNSYQSRADSR | 8 | 1.0785 | 0.37150069 |
| 102233 | 1537 | 785.872314 | 0.007701014 | 2 | TLPFQLSSVEK | 507 | 517 | T | T | TlPFQlSSVEK | 5 | 0.0569 | -1.7108109 |
| 103347 | 1554 | 785.874084 | 6.14E-06 | 2 | TLPFQLSSVEK | 507 | 517 | T | T | TLPfQLSSVEK | 3 | 1.40683 | 0.54148435 |
| 122022 | 1802 | 724.9642 | 8.29E-06 | 3 | VKFDYVPTDTTPR | 826 | 838 | TG-H3N1 | G | VKFDYVpTDTTpR | 11 | 0.22706 | 4.84000531 |
| 9274 | 1409 | 620.955017 | 8.03E-06 | 3 | VKFDYVPTDTTPR | 826 | 838 | T | T | VKfDYVPTDTTPR | 2 | 0.23548 | 4.80471219 |
| 103385 | 1559 | 930.929443 | 7.72E-06 | 2 | VKFDYVPTDTTPR | 826 | 838 | T | T | VKFDyVPTDTTPR | 4 | 0.89512 | 5.40452226 |
| 6571 | 1058 | 556.265686 | 7.99E-06 | 2 | VPLGTFR | 545 | 551 | T | T | VpLGTFR | 1 | 0.09573 | 5.15696366 |
| 7122 | 1128 | 720.791138 | 0.003992608 | 2 | VPLGTFR | 545 | 551 | GT | G | vPLGTFR | 0 | 0.10576 | 2.85707838 |
| 6480 | 1047 | 556.265381 | 7.09E-06 | 2 | VPLGTFR | 545 | 551 | T | T | VPLgTFR | 3 | 0.59214 | 4.60834568 |
| 6341 | 1029 | 556.264709 | 6.78E-06 | 2 | VPLGTFR | 545 | 551 | T | T | VPLgTFR | 3 | 2.40918 | 3.40138614 |

*Mechlorethamine crosslinking sites*

| index | RT | precursor m/z | score | charge | sequence | start position | end position | NuXL:NA | NuXL:NT | NuXL:best_localization | NuXL:best_localization_position | NuXL:best_localization_score | precursor_mz_error_ppm |
|---|---|---|---|---|---|---|---|---|---|---|---|---|---|
| 3472 | 637 | 756.7692 | 6.85E-06 | 2 | CEASEQK | 536 | 542 | AT+C5H9N1 | A | CeASEQK | 1 | 0.1981 | 5.578125 |
| 4255 | 745 | 616.8087 | 5.68E-06 | 2 | VTAVEVGK | 812 | 819 | G+C5H9N1 | G | VTAVeVGK | 4 | 0.8933 | 5.406465 |
| 4754 | 812 | 515.8949 | 7.03E-06 | 3 | VTAVEVGK | 812 | 819 | AG+C5H9N1 | A | VTAVeVGK | 4 | 0.1817 | 5.84546 |
| 5112 | 861 | 510.5626 | 6.96E-06 | 3 | VTAVEVGK | 812 | 819 | AA+C5H9N1 | A | VTaVEVGK | 2 | 0.0576 | 4.539196 |
| 5175 | 870 | 510.5628 | 0.0067337 | 3 | VTAVEVGK | 812 | 819 | AA+C5H9N1 | A | VTAVEVGK | -1 | 0 | 5.017378 |
| 5305 | 886 | 913.0333 | 6.32E-06 | 3 | KDSNELSDSAGEEDSADLK | 771 | 789 | AA+C5H9N1 | A | kDSNELSDSAGEEDSADLK | 0 | 0.1898 | 2.902789 |
| 5553 | 920 | 760.8348 | 5.84E-06 | 2 | VTAVEVGK | 812 | 819 | AT+C5H9N1 | A | VTAVeVGK | 4 | 0.9427 | 5.119838 |
| 5896 | 965 | 717.2875 | 0.0022799 | 2 | TESPIK | 723 | 728 | GG+C5H9N1 | G | TESpIK | 3 | 0.0522 | 4.655995 |

| | | | | | | | | | | | | | |
|---|---|---|---|---|---|---|---|---|---|---|---|---|---|
| 6089 | 990 | 504.9027 | 5.54E-06 | 3 | LSCCLYKPR | 267 | 275 | G+C5H9N1 | G | LScCLYKPR | 2 | 0.2037 | 5.580828 |
| 6178 | 1002 | 504.9029 | 6.24E-06 | 3 | LSCCLYKPR | 267 | 275 | G+C5H9N1 | G | LSCCLYkPR | 6 | 0.0118 | 5.957218 |
| 6336 | 1023 | 504.9029 | 6.07E-06 | 3 | LSCCLYKPR | 267 | 275 | G+C5H9N1 | G | LSCCLYKPr | 8 | 0.181 | 5.943485 |
| 6435 | 1036 | 504.9026 | 6.56E-06 | 3 | LSCCLYKPR | 267 | 275 | G+C5H9N1 | G | LsCCLYKPR | 1 | 0.04 | 5.278614 |
| 6799 | 1084 | 669.2946 | 4.26E-06 | 3 | SNAMAFTNYSSLNR | 0* | 11 | G+C5H9N1 | G | SNAmAFTNYSSLNR | 3 | 1.6757 | 5.321795 |
| 7396 | 1163 | 1118.97 | 5.30E-06 | 2 | SVAVSDEEEVEEEAER | 739 | 754 | G+C5H9N1 | G | SvAvSDEEEvEEEAER | 9 | 0.0075 | 3.685743 |
| 7847 | 1223 | 770.6443 | 4.66E-06 | 3 | SNAMAFTNYSSLNR | 0* | 11 | GT+C5H9N1 | G | SNAmAFTNYSSLNR | 3 | 1.2794 | 5.923861 |
| 9145 | 1391 | 858.6728 | 5.64E-06 | 3 | DYPDTWVCSM(Oxidation)NPDPEQDR | 518 | 535 | C+C5H9N1 | C | DYPDTWVCSMNPDPEQDr | 17 | 0.7754 | 3.468842 |
| 9311 | 1412 | 861.0287 | 6.52E-06 | 3 | QYEVGLQNLCNSYQSR | 953 | 968 | CC+C5H9N1 | C | QYEVGlQNlCNSYQSR | 8 | 0.4286 | 5.299045 |
| 10023 | 1508 | 960.0226 | 6.86E-06 | 3 | DYPDTWVCSM(Oxidation)NPDPEQDR | 518 | 535 | CT+C5H9N1 | C | DYpDTWVCSMNpDpEQDR | 13 | 0.3493 | 4.251615 |
| 10152 | 1525 | 874.3621 | 5.40E-06 | 3 | QYEVGLQNLCNSYQSR | 953 | 968 | CG+C5H9N1 | C | QYEVGLQNLCNsYQSR | 11 | 0.3193 | 2.905539 |
| 10233 | 1536 | 577.2558 | 6.58E-06 | 3 | EDTMTCLFLSR | 122 | 132 | A+C5H9N1 | A | EDTMTcLFLSR | 5 | 0.357 | 5.242686 |
| 10315 | 1547 | 865.379 | 6.56E-06 | 2 | EDTMTCLFLSR | 122 | 132 | A+C5H9N1 | A | eDTMTCLFLSR | 0 | 0.0914 | 3.983867 |
| 10317 | 1547 | 912.7424 | 6.95E-06 | 3 | QYEVGLQNLCNSYQSRADSR | 953 | 972 | T+C5H9N1 | T | QYeVGLQNLCNSYQSRADSR | 2 | 0.2924 | 1.569192 |
| 10401 | 1558 | 865.3795 | 6.42E-06 | 2 | EDTMTCLFLSR | 122 | 132 | A+C5H9N1 | A | EDTMTCLFlSR | 8 | 0.1782 | 4.548109 |
| 10551 | 1577 | 865.3782 | 6.79E-06 | 2 | EDTMTCLFLSR | 122 | 132 | A+C5H9N1 | A | EDtMtCLFLSR | 4 | 0.1786 | 3.137504 |
| 10672 | 1593 | 576.2748 | 0.0067337 | 3 | TNIVALLQK | 999 | 1007 | AA+C5H9N1 | A | TNIVAllQK | 6 | 0.8177 | -5.322833 |
| 10726 | 1600 | 865.3782 | 6.38E-06 | 2 | EDTMTCLFLSR | 122 | 132 | A+C5H9N1 | A | EDTmTCLFLSR | 3 | 0.1091 | 3.066974 |
| 108 | 1611 | 874.0329 | 3.82E-06 | 3 | QYEVGLQNLCNSYQSR | 953 | 968 | AT+C5H9N1 | A | QYEVgLQNLCNSYQSR | 4 | 0.1185 | 5.868916 |

| | | | | | | | | | | | | | |
|---|---|---|---|---|---|---|---|---|---|---|---|---|---|
| 06 | | | | | | | | | | | | | |
| 10866 | 1619 | 865.3795 | 6.80E-06 | 2 | EDTMTCLFLSR | 122 | 132 | A+C5H9N1 | A | EDtMtCLFLSR | 4 | 0.1221 | 4.548109 |
| 10933 | 1628 | 865.3786 | 6.73E-06 | 2 | EDTMTCLFLSR | 122 | 132 | A+C5H9N1 | A | EDtMtCLFLSR | 4 | 0.033 | 3.559444 |
| 11856 | 1747 | 874.0305 | 4.13E-06 | 3 | QYEVGLQNLCNSYQSR | 953 | 968 | AT+C5H9N1 | A | QYEVGLQNLCnSYQSR | 10 | 1.8209 | 3.123007 |
| 12573 | 1839 | 717.9372 | 0.0045219 | 3 | DLDGMFIYNCSR | 378 | 389 | AT+C5H9N1 | A | dLDGMFIYNCSR | 0 | 0.6064 | -4.403546 |
| 12741 | 1861 | 836.3563 | 6.97E-06 | 3 | DLDGM(Oxidation)FIYNCSRLIK | 378 | 392 | AC+C5H9N1 | A | DLDGMfIYNCSRLIK | 5 | 0.8565 | -4.160988 |
| 12758 | 1863 | 874.3637 | 0.0088962 | 3 | QYEVGLQNLCNSYQSR | 953 | 968 | CG+C5H9N1 | C | QYEVgLQNLCNSYQSR | 4 | 0.1792 | 4.780573 |
| 13863 | 2010 | 572.2576 | 0.0067337 | 3 | LLQPPEAPR | 667 | 675 | CT+C5H9N1 | C | LLQppEApR | 7 | 0.0258 | -1.050692 |
| 14557 | 2101 | 1060.118 | 6.18E-06 | 3 | EYFKQYEVGLQNLCNSYQSR | 949 | 968 | TT+C5H9N1 | T | EYFKQYEVGLQNLCNsYQsR | 18 | 0.0814 | 4.167658 |
| 14641 | 2113 | 726.2772 | 6.47E-06 | 3 | DLDGM(Oxidation)FIYNCSR | 378 | 389 | AA+C5H9N1 | A | DLDGMFIyNCSR | 7 | 0.6885 | 1.888239 |
| 14646 | 2113 | 1060.118 | 6.29E-06 | 3 | EYFKQYEVGLQNLCNSYQSR | 949 | 968 | TT+C5H9N1 | T | EYFKQYEVGLQnLCnSYQSR | 14 | 0.1186 | 4.450646 |
| 14996 | 2160 | 942.938 | 0.0045219 | 2 | PANTLVKTASR | 677 | 687 | AA+C5H9N1 | A | PANTlVKTASR | 4 | 0.1258 | 3.044834 |
| 15086 | 2171 | 942.9363 | 0.0088962 | 2 | PANTLVKTASR | 677 | 687 | AA+C5H9N1 | A | PANTlVKTASR | 4 | 0.1371 | 1.219997 |
| 15271 | 2196 | 942.9353 | 7.01E-06 | 2 | PANTLVKTASR | 677 | 687 | AA+C5H9N1 | A | PANTlVKTASR | 4 | 0.1415 | 0.181435 |
| 15538 | 2231 | 700.8522 | 7.00E-06 | 2 | EYRHLLR | 435 | 441 | A+C5H9N1 | A | EYRHLlR | 5 | 1.0685 | 1.777376 |

*The start position of this peptide includes a three amino acid scar from the cleavage of the affinity tag.

**SI Table 4. $V_{max}$ and $K_m^{app,ATP}$ of MORC2 in the presence of various DNA concentrations.**

| | **0µM DNA** | **0.05µM DNA** | **0.1µM DNA** | **1µM DNA** |
|---|---|---|---|---|
| $V_{max}$ ($min^{-1}$) | 0.41 ± 0.02 | 0.36 ± 0.03 | 0.22 ± 0.02 | 0.17 ± 0.01 |
| $K_m^{app, ATP}$ (µM) | 0.4 ± 0.1 | 0.5 ± 0.1 | 0.4 ± 0.1 | 0.4 ± 0.1 |

**SI Figure 1. Assessment of MORC2 protein and substrate purity.**

**A.** SDS-PAGE gel of purified MORC2 proteins stained with Coomassie blue stain. Gel was run with 10 μg of each protein to show the level of purity after size-exclusion chromatography.

**B.** Circular dichroism spectrometry of wildtype and mutant MORC2. Spectra were taken at 25˚C (**Methods**).

**C.** Native PAGE gel of reconstituted nucleosome stained with SYBR gold.

**SI Figure 2. Representative spectra from mechlorethamine and UV-induced protein-DNA crosslinking mass spectrometry.**

The relative intensity of MS/MS ions is plotted against their mass-to-charge ratios (*m/z*). Red corresponds to a and b ions, and green corresponds to y ions of the peptide. The sequence of the identified peptide is displayed with the detected ions colored in red or green. Spectra shown from **(A)** peptide fragment K722-K728 crosslinked by UV to dTMP, (**B**) peptide fragment G762-K767 crosslinked by UV to dCMP, (**C**) peptide fragment T723-K728 crosslinked by mechlorethamine to dGMP, and (**D**) peptide fragment S739-R754 crosslinked by mechlorethamine to dGMP. Spectra were exported from TOPPView (**Methods**) and for (**C**) the precursor ion ([M+2H] $NM+GG^{2+}$) was manually annotated. NM = nitrogen mustard or mechlorethamine.

**SI Figure 3. Sequence alignment of MORC2 C-terminal domain.**

MORC2 sequences from the indicated organisms were aligned in MAFFT and visualized in Jalview. Residues colored by percentage identity. Darker shades of blue indicate higher conservation. Phosphorylation sites are indicated in orange and putative DNA binding residues are indicated in pink.

**SI Figure 4. ATPase activity and DNA binding of MORC2 mutants.**

**A.** Assessment of MORC2 ATPase activity. Phosphomimetic, subset N, subset C, alanine mutant, and phosphodead MORC2 (1 μM) were incubated with 1 mM ATP for 45 minutes at 37°C either in the presence or absence of 2 μM of a 35 base pair dsDNA. Inorganic phosphate released was quantified by malachite green (Methods). Error bars correspond to the standard deviation between three replicate experiments.

**B.** Assessment of DNA binding by alanine mutant, subset N, and subset C MORC2. MORC2 constructs were titrated and incubated with 1 nM FAM-labelled of a 35 base pair duplex DNA (Methods). Data were fit using a quadratic binding equation. Error bars correspond to the standard deviation between three replicate experiments.

**C.** MORC2 DNA binding to DNA sequences of different lengths as assessed by fluorescence anisotropy. MORC2 was titrated and incubated with 1 nM 5’ FAM-labelled 25 base pair duplex DNA, 45 base pair duplex DNA, and 65 base pair duplex DNA (Methods). Data were fit using a quadratic binding equation. Error bars correspond to the standard deviation between three replicate experiments.

**D.** MORC2 DNA binding in the presence of ATP analogs. MORC2 was titrated and incubated with 1 nM 5’ FAM-labelled 35 base pair duplex DNA in the presence or absence of 1 mM AMP-PNP, ATPγS, ADP, or ATP (**Methods**). Data were fit using a quadratic binding equation. Error bars correspond to the standard deviation between three replicate experiments.

**SI Figure 5. Dimerization interfaces of MORC2.**

AlphaFold Multimer model of full-length MORC2 as a dimer. Chain A is colored grey and chain B is colored by pLDDT score, as shown below the model. The end of the crystal structure model is indicated. Highlighted in magenta is the region of the C-terminus encompassing the phosphorylation sites and positively charged residues mutated.

**SI Figure 6. Replicate gels from Figure 4.**

**A.** N-terminal-maltose binding protein (MBP)-tagged wildtype, aspartate mutant, and E35A MORC2 (600 nM) were incubated with 100 nM supercoiled or linear pUC19. Samples were then added to amylose resin and washed in either low salt (50 mM NaCl) or high salt (400 mM NaCl) containing buffers before eluting the samples from the beads with maltose. Eluted samples were treated with proteinase K. DNA was resolved on a 1% (w/v) TAE agarose gel.

**B.** A biotin tagged DNA was conjugated to streptavidin magnetic beads to create a pseudo circular substrate. MORC2 (600 nM) was incubated with 20 μL of the beads in the presence or absence of 1 mM AMP-PNP. Supercoiled pBlueScript plasmid DNA (200 nM) was added before washing the beads with either low salt (50 mM NaCl) or high salt (400 mM NaCl) containing buffer. Samples were resuspended in 1X CutSmart buffer (New England Biolabs). DNA was released from the beads by digestion with ScaI and SbfI at 37°C for 1 hour before proteinase K treatment. DNA was resolved on a 1% (w/v) TAE agarose gel.

**SI Figure 7. MORC2 localization analysis.**

**A.** NLS Stradmus analysis of potential nuclear localization sequences in MORC2 (**Methods**). Positively charged residues mutated in this study are shown below the graph in pink.

**B.** Representative confocal microscopy images of interphase HeLa cells overexpressing EGFP- subset N, EGFP- E35A, EGFP- N39A, and EGFP- aspartate mutant MORC2.

**C.** Representative confocal microscopy images of interphase HeLa cells overexpressing artificial NLS-EGFP, artificial NLS-EGFP- wildtype, and artificial NLS-EGFP- aspartate mutant MORC2.

**SI Figure 8. MORC2 knockout validation.**

**A.** Sanger sequencing of MORC2 gene locus in parental and knockout HeLa cells.

**B.** PCR amplification with primers designed to amplify a region of the MORC2 gene locus encompassing the deleted sequence of parental and knockout (KO) HeLa cell genomic DNA. DNA products were separated on a 1% TAE agarose gel and stained with SYBR safe. The PCR product of the native locus is 1199 bp.

**C.** Western blot analysis of MORC2 protein level in parental and knockout (KO) HeLa cells. Beta-actin was used as a loading control.

**SI Figure 9. RNA sequencing controls and analysis.**

**A.** Genome browser trace of MORC2 gene locus from NLS-EGFP (control), NLS-aspartate mutant, and NLS-wildtype MORC2 samples with exon 19, which contains the positive charge residue mutations in the aspartate mutant.

**B.** Comparison of exogenous MORC2 RNA levels in NLS-EGFP (control), NLS-aspartate mutant, and NLS-wildtype MORC2 samples.

**C.** Principle component analysis of the three biological replicates of NLS-EGFP, NLS-aspartate mutant, and NLS-wildtype MORC2 samples.

**D.** Volcano plots of RNAseq reads for overexpression of wildtype MORC2 versus control, overexpression of aspartate mutant MORC2 versus control, and wildtype MORC2 versus aspartate mutant MORC2 without spike normalization (**Methods**). Significant upregulated genes are shown in green and significant downregulated genes are shown in purple from three biological replicates. Significant genes are classified as those that meet the fold change > 1.5 and FDR > 0.05 cutoffs.

**E.** Venn diagram representation of the overlap between the significantly downregulated genes after overexpression of wildtype MORC2 identified in this study and previously identified targets of MORC2 silencing.

**F.** Representation of significant downregulated genes after overexpression of wildtype MORC2 in HeLa cells. 75 out of 197 genes are intronless or contain exons longer than 1 kb.

### SI Fig 1

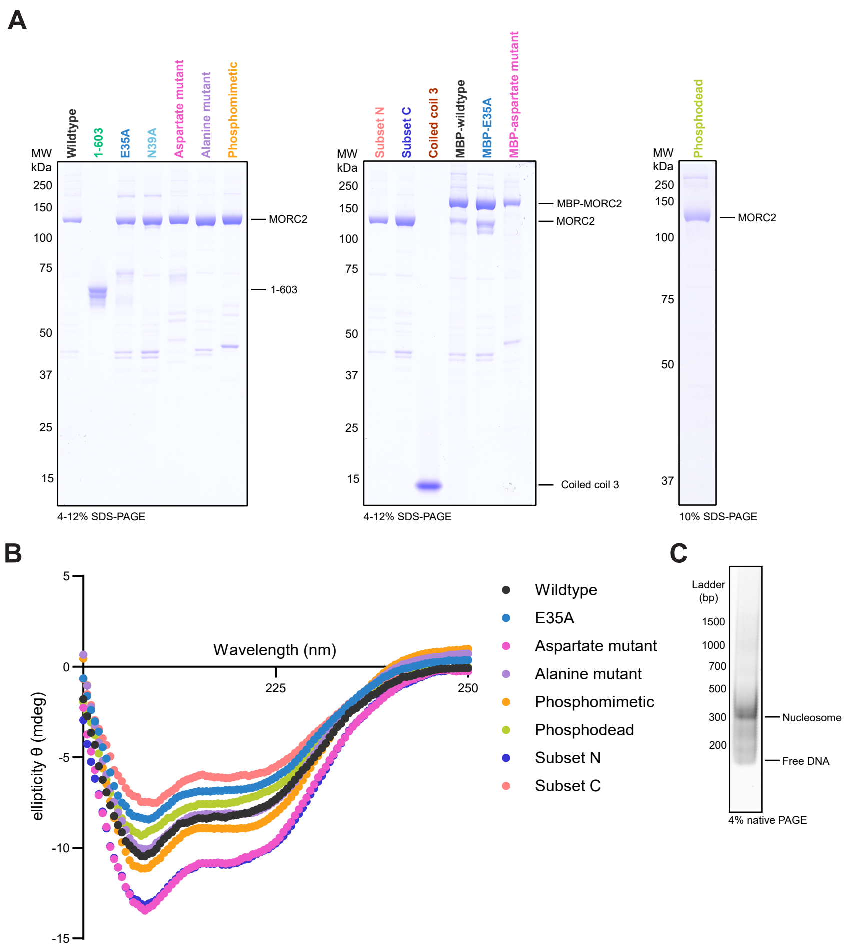
A
MW kDa
Wildtype
1-603
E35A
N39A
Aspartate mutant
Alanine mutant
Phosphomimetic
250
150
100
75
50
37
25
15
MORC2
1-603
4-12% SDS-PAGE
Subset N
Subset C
Coiled coil 3
MBP-wildtype
MBP-E35A
MBP-aspartate mutant
MBP-MORC2
MORC2
Coiled coil 3
4-12% SDS-PAGE
Phosphodead
MORC2
10% SDS-PAGE
B
Wavelength (nm)
225
250
ellipticity θ (mdeg)
5
0
-5
-10
-15
Wildtype
E35A
Aspartate mutant
Alanine mutant
Phosphomimetic
Phosphodead
Subset N
Subset C
C
Ladder (bp)
1500
1000
700
500
300
200
Nucleosome
Free DNA
4% native PAGE

#### SI Fig 2

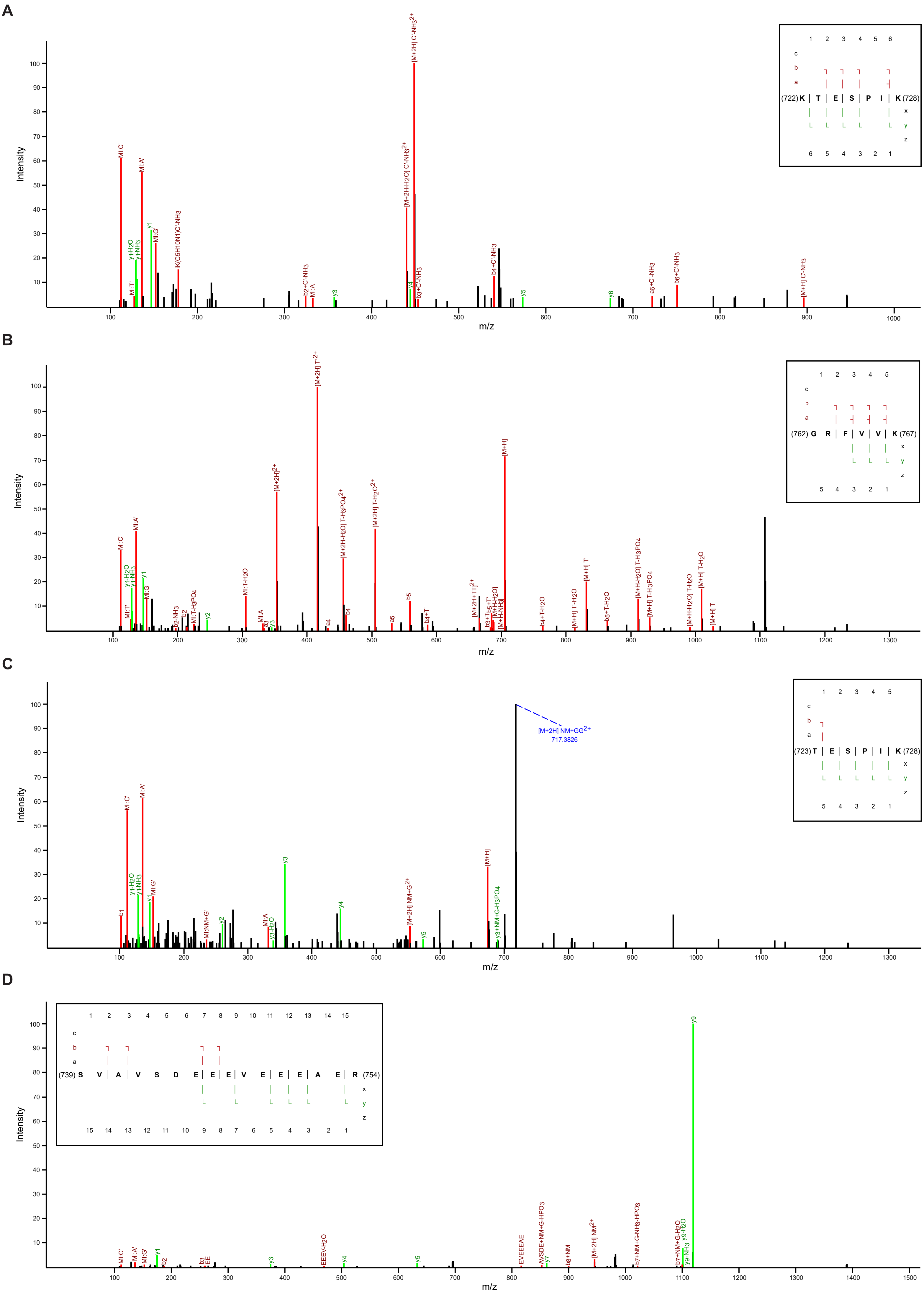

A
(722) K T E S P I K (728)
Intensity
m/z
[M+2H] C'-NH3 2+
[M+2H-H2O] C'-NH3 2+
MI:C'
MI:A'
y1
MI:G'
y-H2O
y-NH3
MI:T'
K(C5H10N1)C'-NH3
b2+C'-NH3
MI:A
y3
y4
b3+C'-NH3
b4+C'-NH3
y5
y6
a6+C'-NH3
b6+C'-NH3
[M+H] C'-NH3
B
(762) G R F V V K (767)
Intensity
m/z
[M+2H] T' 2+
[M+H]
[M+2H] 2+
[M+2H] T-H2O 2+
[M+2H-H2O] T-H3PO4 2+
MI:C'
MI:A'
y1-H2O
y1-NH3
y1
MI:G'
MI:T'
b2-NH3
b2
MI:T-H3PO4
y2
MI:T-H2O
MI:A
a3
y3
a4
b4
a5
b5
b4+T'
[M+H] T'
[M+H] T-H2O
[M+H] T'-H2O
[M+H+H2O] T-H3PO4
[M+H] T-H3PO4
[M+H+H2O] T-H2O
[M+H] T
b4+T-H2O
b5+T-H2O
C
(723) T E S P I K (728)
Intensity
m/z
[M+2H] NM+GG 2+
717.3826
MI:C'
MI:A'
y1-H2O
y1-NH3
y1
MI:G'
b1
MI:NM+G'
y2
MI:A
y3-H2O
y3
y4
[M+2H] NM+G 2+
y5
[M+H]
y3+NM+G-H3PO4
D
(739) S V A V S D E E E V E E E A E R (754)
Intensity
m/z
y9
MI:C'
MI:A'
MI:G'
y1
b2
b3
EE
y3
EEEV-H2O
y4
y5
EVEEEAE
AVSDE+NM+G-HPO3
y7
b8+NM
[M+2H] NM 2+
b7+NM+G-NH3-HPO3
b7+NM+G-H2O
y9-NH3
y9-H2O

### SI Fig 3

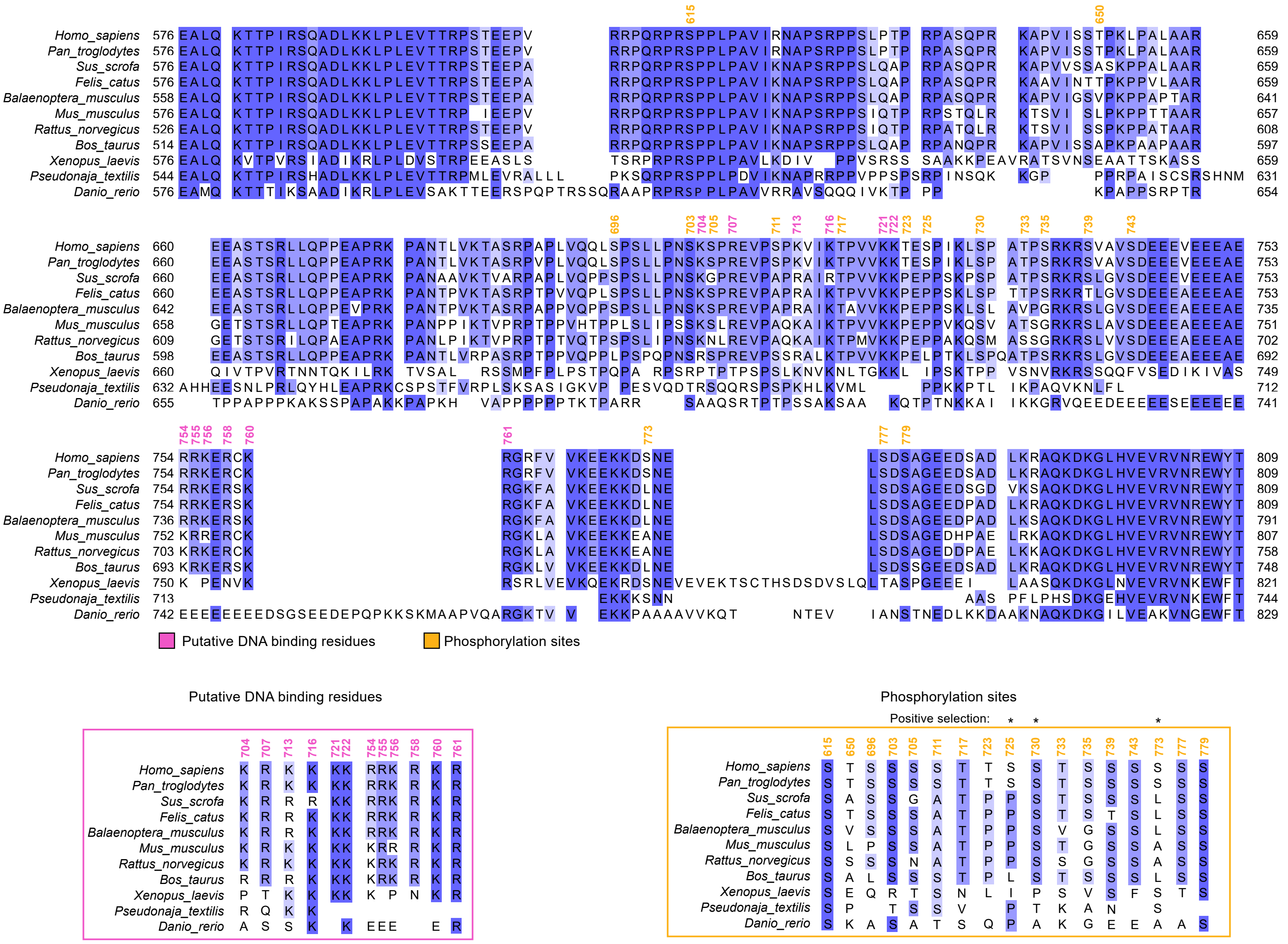
Homo_sapiens
Pan_troglodytes
Sus_scrofa
Felis_catus
Balaenoptera_musculus
Mus_musculus
Rattus_norvegicus
Bos_taurus
Xenopus_laevis
Pseudonaja_textilis
Danio_rerio
Putative DNA binding residues
Phosphorylation sites
Putative DNA binding residues
Phosphorylation sites
Positive selection:

SI Fig 4

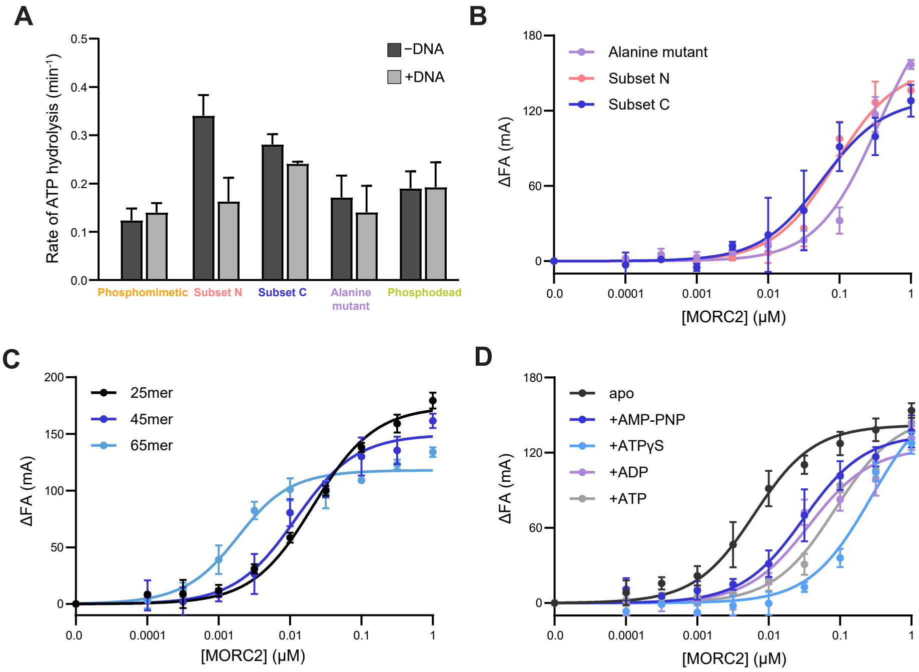
A
B
C
D
Rate of ATP hydrolysis (min⁻¹)
−DNA
+DNA
Phosphomimetic
Subset N
Subset C
Alanine mutant
Phosphodead
ΔFA (mA)
[MORC2] (μM)
Alanine mutant
Subset N
Subset C
25mer
45mer
65mer
apo
+AMP-PNP
+ATPγS
+ADP
+ATP

**SI Fig 5**

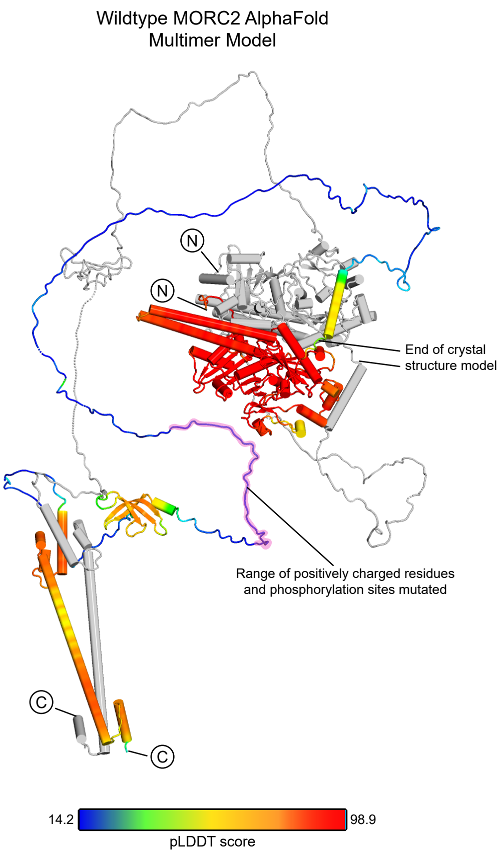

Wildtype MORC2 AlphaFold
Multimer Model
N
N
End of crystal
structure model
Range of positively charged residues
and phosphorylation sites mutated
C
C
14.2
98.9
pLDDT score

### SI Fig 6

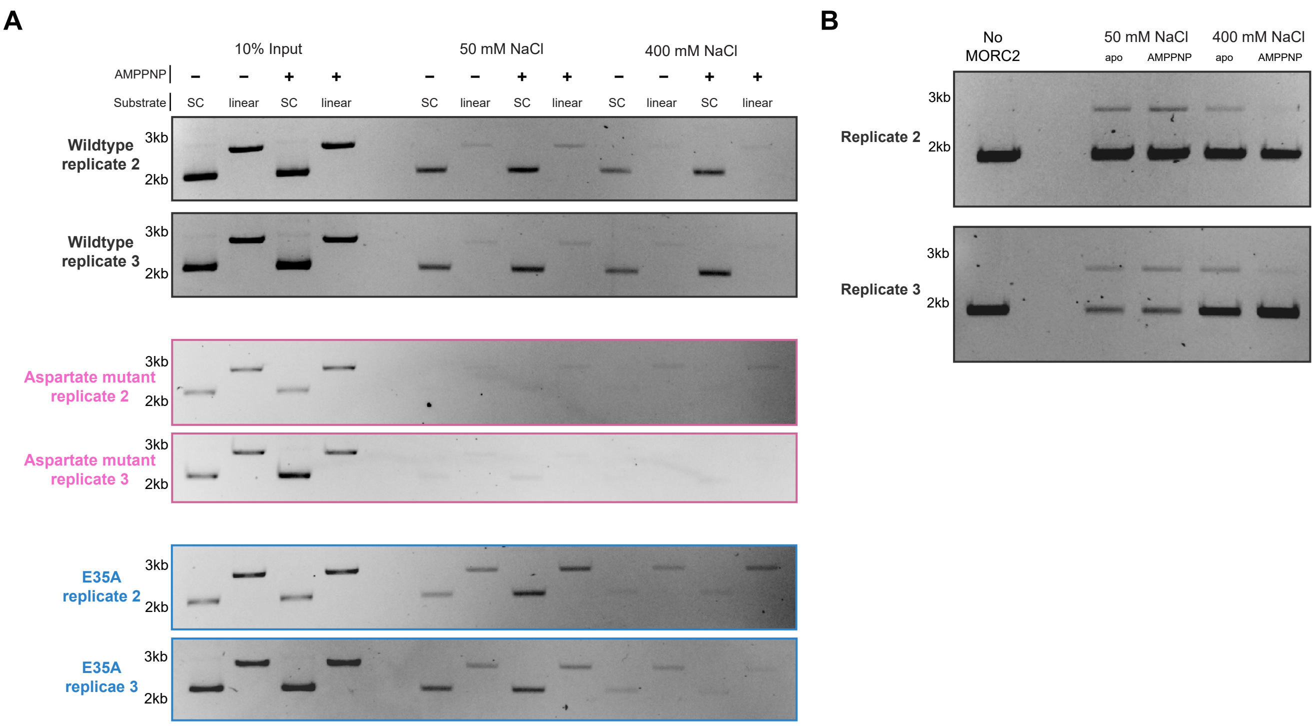

A
10% Input
50 mM NaCl
400 mM NaCl
AMPPNP
Substrate
SC
linear
3kb
2kb
Wildtype replicate 2
Wildtype replicate 3
Aspartate mutant replicate 2
Aspartate mutant replicate 3
E35A replicate 2
E35A replicae 3
B
No MORC2
apo
AMPPNP
Replicate 2
Replicate 3

SI Fig 7

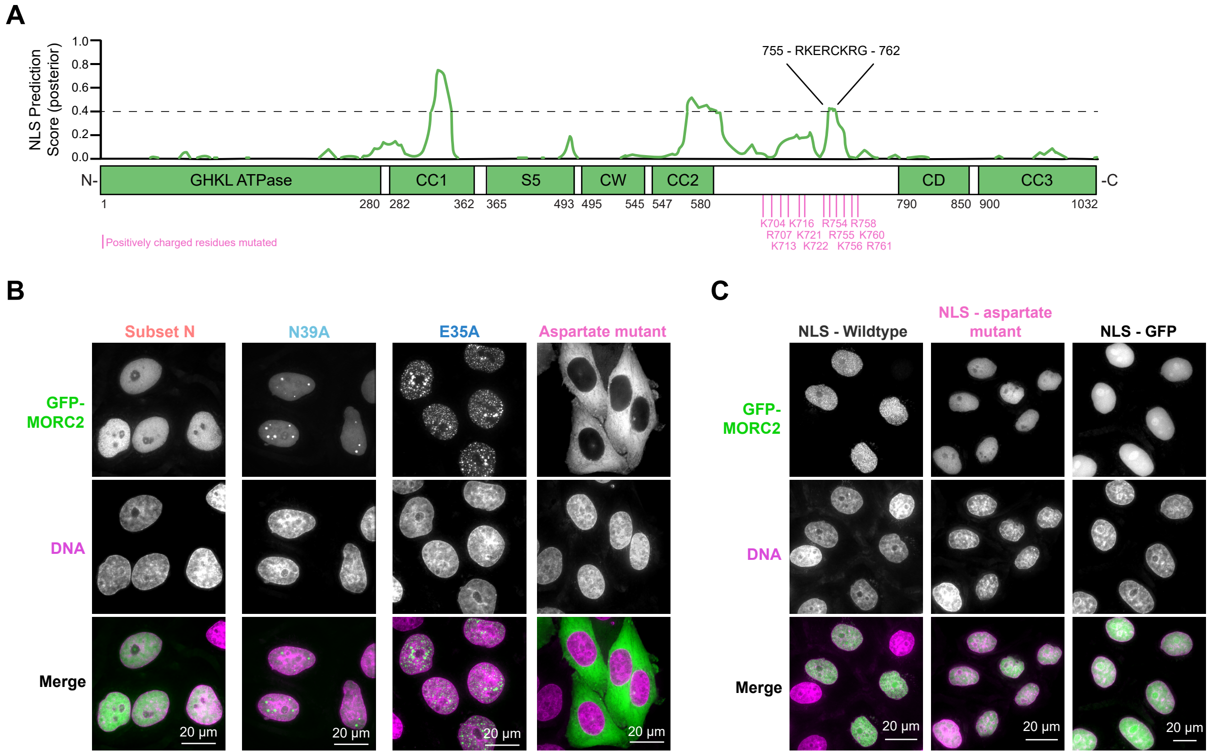
A
NLS Prediction Score (posterior)
1.0
0.8
0.6
0.4
0.2
0.0
755 - RKERCKRG - 762
N-
GHKL ATPase
CC1
S5
CW
CC2
CD
CC3
-C
1
280
282
362
365
493
495
545
547
580
790
850
900
1032
K704 K716 R754 R758
R707 K721 R755 K760
K713 K722 K756 R761
Positively charged residues mutated
B
Subset N
N39A
E35A
Aspartate mutant
GFP-MORC2
DNA
Merge
20 µm
C
NLS - Wildtype
NLS - aspartate mutant
NLS - GFP

### SI Fig 8

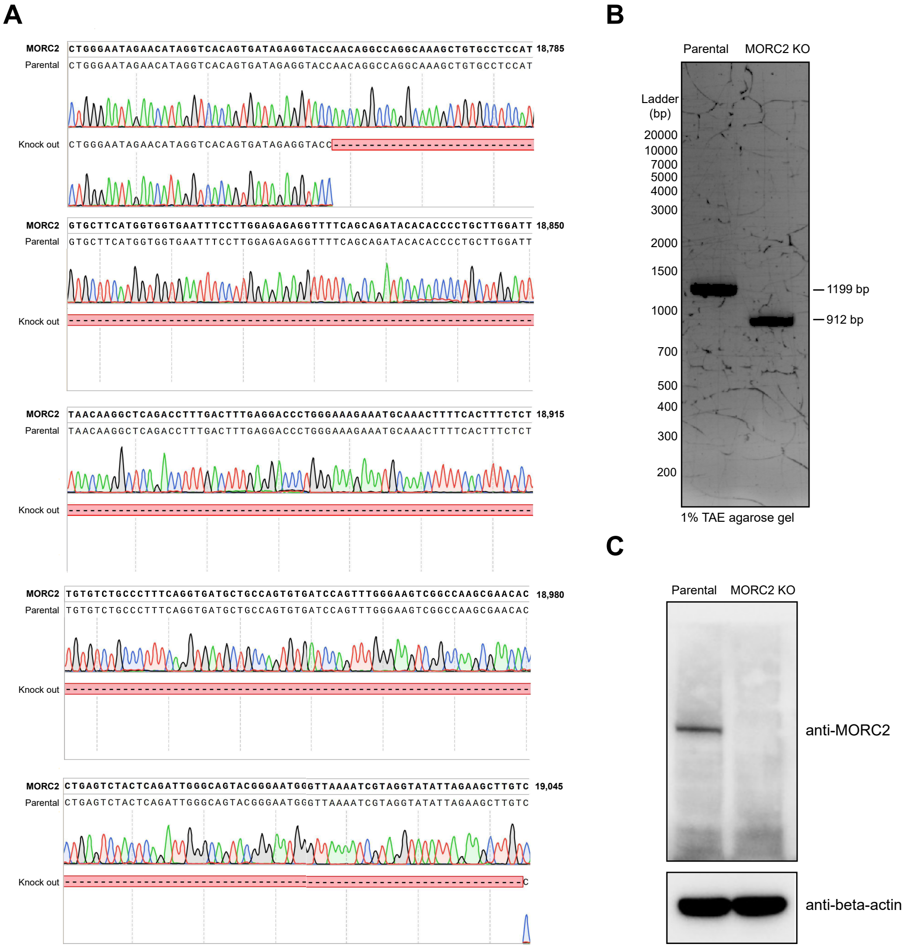

A
MORC2 CTGGGAATAGAACATAGGTCACAGTGATAGAGGTACCAACAGGCCAGGCAAAGCTGTGCCTCCAT 18,785
Parental CTGGGAATAGAACATAGGTCACAGTGATAGAGGTACCAACAGGCCAGGCAAAGCTGTGCCTCCAT
Knock out CTGGGAATAGAACATAGGTCACAGTGATAGAGGTACC
MORC2 GTGCTTCATGGTGGTGAATTTCCTTGGAGAGAGGTTTTCAGCAGATACACACCCCTGCTTGGATT 18,850
Parental GTGCTTCATGGTGGTGAATTTCCTTGGAGAGAGGTTTTCAGCAGATACACACCCCTGCTTGGATT
Knock out
MORC2 TAACAAGGCTCAGACCTTTGACTTTGAGGACCCTGGGAAAGAAATGCAAACTTTTCACTTTCTCT 18,915
Parental TAACAAGGCTCAGACCTTTGACTTTGAGGACCCTGGGAAAGAAATGCAAACTTTTCACTTTCTCT
Knock out
MORC2 TGTGTCTGCCCTTTCAGGTGATGCTGCCAGTGTGATCCAGTTTGGGAAGTCGGCCAAGCGAACAC 18,980
Parental TGTGTCTGCCCTTTCAGGTGATGCTGCCAGTGTGATCCAGTTTGGGAAGTCGGCCAAGCGAACAC
Knock out
MORC2 CTGAGTCTACTCAGATTGGGCAGTACGGGAATGGGTTAAAATCGTAGGTATATTAGAAGCTTGTC 19,045
Parental CTGAGTCTACTCAGATTGGGCAGTACGGGAATGGGTTAAAATCGTAGGTATATTAGAAGCTTGTC
Knock out C
B
Parental
MORC2 KO
Ladder (bp)
20000
10000
7000
5000
4000
3000
2000
1500
1000
700
500
400
300
200
1199 bp
912 bp
1% TAE agarose gel
C
Parental
MORC2 KO
anti-MORC2
anti-beta-actin

#### SI Fig 9

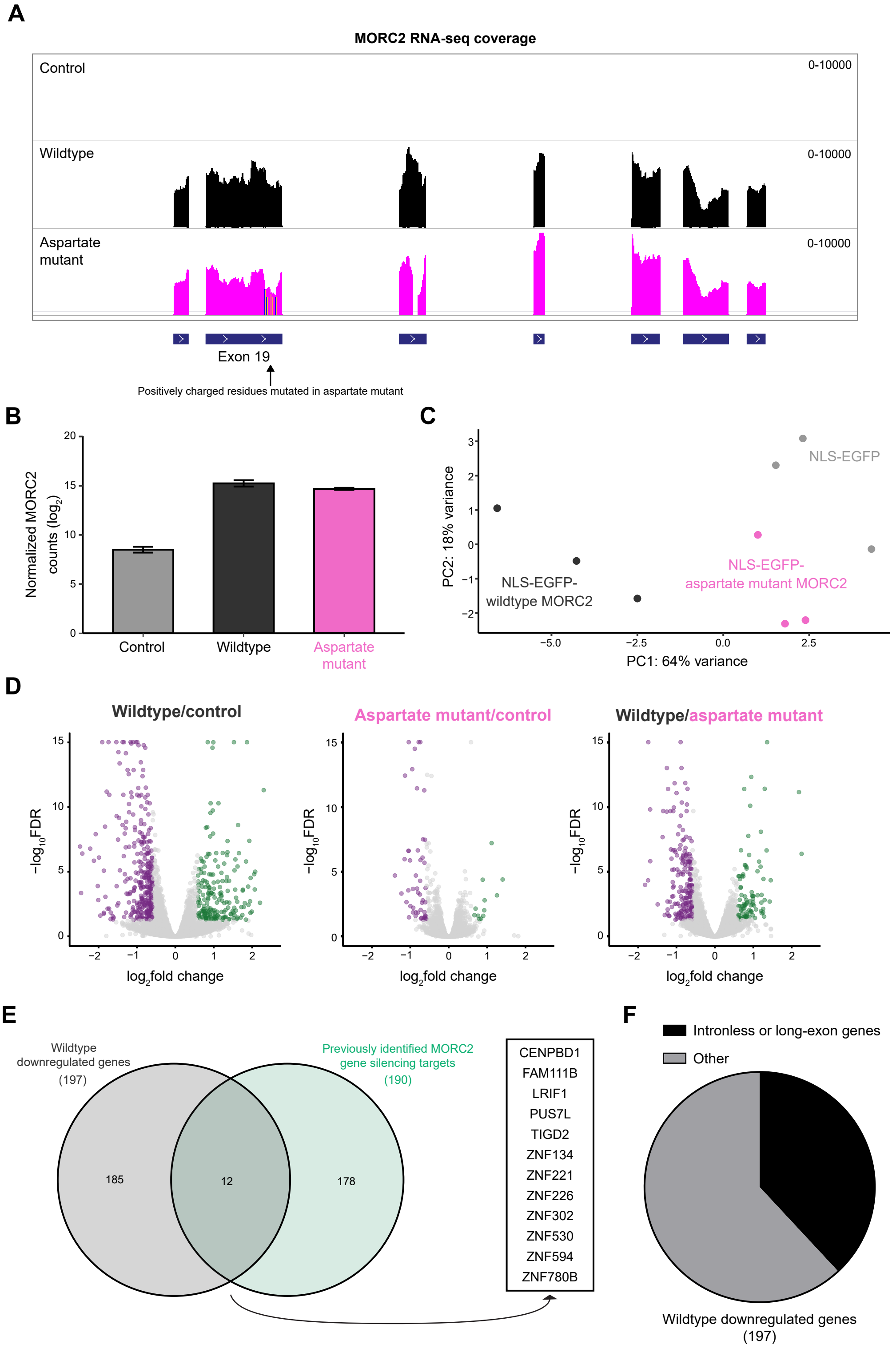
A
MORC2 RNA-seq coverage
Control
0-10000
Wildtype
0-10000
Aspartate mutant
0-10000
Exon 19
Positively charged residues mutated in aspartate mutant
B
Normalized MORC2 counts (log2)
Control
Wildtype
Aspartate mutant
C
PC2: 18% variance
PC1: 64% variance
NLS-EGFP
NLS-EGFP-wildtype MORC2
NLS-EGFP-aspartate mutant MORC2
D
Wildtype/control
Aspartate mutant/control
Wildtype/aspartate mutant
−log10FDR
log2fold change
E
Wildtype downregulated genes (197)
Previously identified MORC2 gene silencing targets (190)
185
12
178
CENPBD1
FAM111B
LRIF1
PUS7L
TIGD2
ZNF134
ZNF221
ZNF226
ZNF302
ZNF530
ZNF594
ZNF780B
F
Intronless or long-exon genes
Other
Wildtype downregulated genes (197)
